# The microcephaly-associated protein YIPF5 differentially regulates ER-export

**DOI:** 10.1101/2025.06.11.659036

**Authors:** Francesca Bruno, Mihaela Anitei, Domenico Di Fraia, William Durso, Therese Dau, Emilio Cirri, Mara Sannai, Christina Valkova, Julija Maldutyte, Elizabeth A. Miller, Ignacio Rubio, Vera Garloff, Noortje Kersten, Ginny Farias, Alessandro Ori, Iván Mestres, Federico Calegari, Christoph Kaether

## Abstract

YIPF5 is a small ER-membrane protein implicated in ER-Golgi transport. Mutations in YIPF5 cause MEDS2 (microcephaly with simplified gyral pattern, epilepsy, and neonatal diabetes syndrome), a fatal disorder manifesting in early childhood. We demonstrate that YIPF5 is involved in ER export of a subset of proteins, including cargoes of the ER-export receptor SURF4, with which it directly interacts. In YIPF5 knockout cells, we observe a shift in the cell surface and secretome composition, marked by reduced neuronal adhesion molecules and increased secretion of ER chaperones influencing cell migration. YIPF5 depletion enhances cell migration in a wound-healing assay and alters SURF4 localization, causing elongated ERGIC53- and Rab1-positive tubules from COPII-labeled ER exit sites. Kinetic analysis suggests that YIPF5 negatively regulates SURF4-mediated ER export. *In utero* knockdown of *Yipf5* in embryonic mouse brains induces premature neuronal migration and abnormal neuronal morphology. These findings suggest that YIPF5 and SURF4 coordinate ER export, and disruption may underlie cortical development defects leading to microcephaly.

## Introduction

Membrane and secreted proteins, roughly representing one third of all proteins, are mostly synthesized at the endoplasmic reticulum (ER), the largest organelle in the cell. Proteins destined for downstream locations along the secretory pathway are exported from the ER at specific sites, termed ER-exit sites (ERES) (Bannykh et al., 1996; Hammond and Glick, 2000; Kurokawa and Nakano, 2019). Here, a sophisticated machinery consisting of the membrane-bending small GTPase Sar1 and the COPII-coat protein complexes SEC23/SEC24 and SEC13/SEC31 concentrate and pack cargo proteins into ER-Golgi transport carriers (Aridor, 2022; Barlowe et al., 1994). The exact role of COPII in ER export is controversial, with recent studies suggesting that instead of coating small vesicles, the COPII-coat is localized at the neck of forming transport carriers and actually never leaves the ERES (Shomron et al., 2021; Weigel et al., 2021), reviewed in (Downes and Zanetti, 2025; Malhotra, 2024; Malis et al., 2022; Tang and Ginsburg, 2023). The basic COPII machinery is sufficient to mediate ER export *in vitro*. To concentrate cargo, in cells it is assisted by a number of adaptors and helper proteins like the p24 family of adaptors, cornichons and SURF4 (Barlowe and Helenius, 2016; Zhang et al., 2023). SURF4 (surfeit locus protein 4, (Williams et al., 1988)) is a 30 kDa protein with seven predicted transmembrane domains and a C-terminal KKXX ER-retrieval motif (Reeves and Fried, 1995). SURF4 is an ER-Golgi cargo receptor for a subset of cargoes like proinsulin (Saegusa et al., 2022), STING (Deng et al., 2020; Mukai et al., 2021), PCSK9, Cab45 and NUCB1 (Maldutyte et al., 2025) and others (reviewed in (Shen et al., 2023)).

In yeast, COPII-mediated ER-Golgi transport requires the Yip1p-Yif1p-Yos1p complex (Heidtman et al., 2005). The mammalian orthologue of Yip1p, YIPF5 (Yip1 domain family protein 5, aliases FinGER5, YIP1A, YIP1) (Tang et al., 2001) is a member of the YIPF family proteins, consisting of YIPF1-7 (Shaik et al., 2019). Shaik et al. proposed a novel nomenclature for YIPF proteins and suggested YIPFα1A for YIPF5 (Shaik et al., 2019). YIPF5 is a small five-span transmembrane domain (TMD) protein of the ER and the early secretory pathway. It interacts with SEC23/24 (Tang et al., 2001) and in yeast is involved in COPII vesicle biogenesis (Heidtman et al., 2003). YIPF5 is necessary for ER export of the innate immune receptor STING (Ran et al., 2019), is critically involved in the activation of IRE1 (Taguchi et al., 2017; Taguchi et al., 2015) and has been implicated in ER structure (Dykstra et al., 2010). Mutations in *YIPF5* can cause MEDS2 (microcephaly, epilepsy, and diabetes syndrome, OMIM # 614231), a severe syndrome leading to death in early childhood (De Franco et al., 2020). To date, four missense mutations p.(Ala181Val), p.(Ile98Ser), p.(Trp218Arg), p.(Gly97Val) and one in-frame deletion p.(Lys106del) were identified as causative for MEDS2 (De Franco et al., 2020). Loss or mutation of YIPF5 results in proinsulin retention in the ER and increased sensitivity to ER stress, resulting in apoptosis of β-cells (De Franco et al., 2020). One mutation, p.(Trp218Arg), caused microcephaly in rabbits (Liu et al., 2023). IER3IP1, the mammalian orthologue of Yos1p, a second component of the Yip1p-Yif1p-Yos1p complex, is also a small ER-Golgi membrane protein with two predicted TMDs (Yiu et al., 2004). Interestingly, mutations in *IERIP1* cause MEDS1 (Abdel-Salam et al., 2012; Poulton et al., 2011; Shalev et al., 2014). IER3IP1 is involved in the ER-export of a subset of proteins, often involved in neurodevelopmental processes like migration, cell-cell contact and axon pathfinding (Anitei et al., 2024; Esk et al., 2020). IER3IP1 also controls KDELR2 localization, resulting in an enhanced secretion of KDEL-containing ER-resident proteins in its (Anitei et al., 2024). MEDS1 and MEDS2 are characterized by similar phenotypes, supporting the notion that YIPF5 and IER3IP1 interact and have similar molecular functions. Mutations in the gene of the mammalian orthologue of the third complex component Yif1p, *YIF1B*, can also cause microcephaly and epilepsy (AlMuhaizea et al., 2020; Medico Salsench et al., 2021), further suggesting that the YIPF5-IER3IP1-YIF1B complex plays an important role in neurodevelopment.

We here show that YIPF5 interacts with SURF4, controlling ER-export of a specific subset of proteins. In the absence of YIPF5, SURF4-specific cargoes and KDEL-carrying ER resident proteins are enriched in the supernatant, whereas neuronal cell adhesion molecules are reduced at the plasma membrane. Knock-down of *YIPF5* in cultured cells results in faster migration in wound healing assays. Knock-down of mouse *Yipf5* by *in-utero* electroporation at embryonic day E13.5 mice induced over-migration of cortical neuronal precursors and newborn neurons or premature differentiation, providing a mechanism for the microcephaly observed in MEDS patients.

## Results

### YIPF5 localizes to the early secretory pathway, is dispensable for cell growth, but is essential for ER morphology

To study the function of YIPF5, CRISPR-Cas9 was utilized to knockout *YIPF5* in MCF10A cells by directing sgRNA towards exon 3 (Cramer et al., 2023). Additionally, we stably re-expressed YIPF5 wildtype (wt) and the pathogenic variants I98S, L108del, A181V or W218R in YIPF5-KO cells by lentiviral transduction (Fig. 1a). We validated the deletion of YIPF5 and the re-expression of YIPF5 variants in YIPF5-KO cells via immunofluorescence and immunoblotting using a specific antibody against YIPF5 (Fig 1a-c). We observed that endogenous YIPF5 localized to the early secretory pathway, where it partially colocalized with ERGIC-53, a marker of the ER-Golgi intermediate compartment (ERGIC, Suppl. Fig. 1a), and with GM130, a marker of the cis-Golgi apparatus (GM130, Suppl. Fig. 1b), as well as with SEC31A, a marker of ER exit sites (ERES) (Fig. 1a). Notably, the disease-associated I98S, L108del and A181V mutants exhibited a subcellular distribution similar to that of endogenous and re-expressed wild-type YIPF5 (Fig. 1a). In contrast, the W218R mutant predominantly localized to the ER. The absence of YIPF5 resulted in a 2-fold increase in the number of ERES/cell that was normalized by re-expression of YIPF5 wt and the pathogenic mutations except YIPF5 I98S (Fig. 1b). YIPF5 I98S was therefore selected for further investigation.

**Figure 1.**
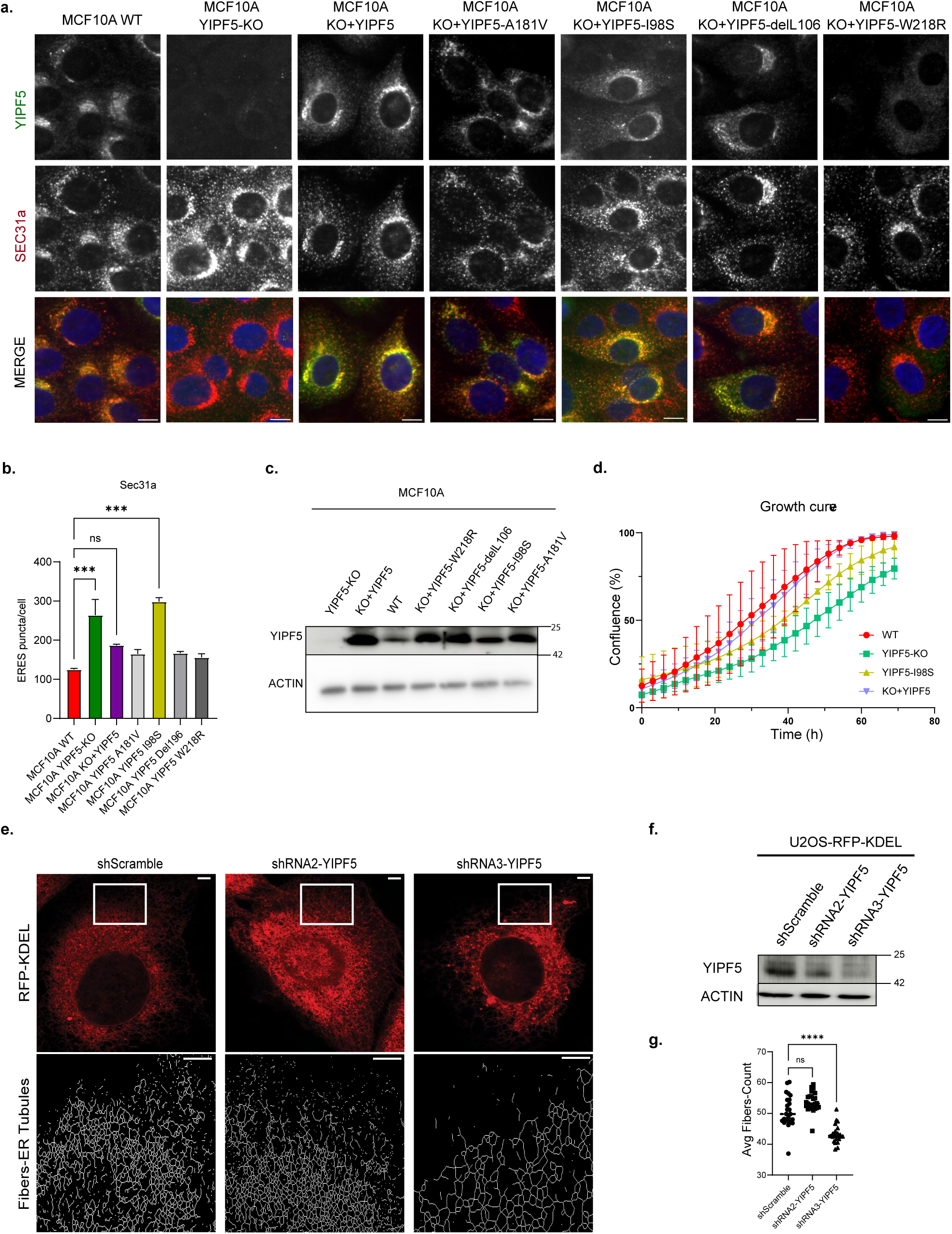
YIPF5 localizes to the early secretory pathway and is dispensable for cell proliferation. (a) MCF10A, MCF10A YIPF5-KO, and MCF10A YIPF5-KO re-expressing YIPF5 variants were fixed and co-labeled with anti-YIPF5 (green) and anti-SEC31A antibodies (red). Nuclei were stained with Hoechst dye (blue). Cells were analyzed by ImageXpress High-content confocal fluorescence microscopy. Scale bars: 10 µm. (b) Image analysis was performed using an algorithm created in the MetaXpress Custom Module Editor Pipeline, developed with MetaXpress software. Displayed is the average number of ERES/cell (n = 3; number of cells analyzed: WT 2089, YIPF5-KO 2325, KO+YIPF5 2699, KO+YIPF5 A181V 1628, KO+YIPF5 I98S 2256, KO+YIPF5-del105 1456, KO+YIPF5 W218R 2549, One-way ANOVA with Tukey’s multiple comparisons test, *** indicates p < 0.001). (c) Western blot analysis of MCF10A cell lines using anti-YIPF5 and anti-Actin (loading control). A representative blot of n = 3 independent experiments is shown. For full-size blots see Suppl. Fig. 6. (d) Incucyte growth curve analysis of MCF10A WT, MCF10A YIPF5-KO, and MCF10A YIPF5-KO re-expressing YIPF5 WT or I98S mutant. Cell growth displayed as % confluence was recorded every 3 h for 72 h. Data were normalized to t0-Control. Data are presented as mean ± SD. (e) Top: U2OS-RFP-KDEL cells stably expressing the indicated shRNAs were imaged by ZEISS Airyscan super-resolution confocal microscopy. Bottom: Application of the ImageJ “skeletonize” function to the boxed regions (top panels) allowed visualization of ER network morphology. Scale bars: top = 10 µM, bottom = 2 µM. (f) Validation of YIPF5 knockdown in U2OS-RFP-KDEL cell lines by Western blot using anti-YIPF5 antibody. Anti-Tubulin served as the loading control. A representative blot of n = 3 independent experiments is shown. Source data are available for this figure: SourceData F1. (g) Quantification of live fluorescence microscopy and high-content confocal fluorescence microscopy analysis of cells from e) plated on a 96-well glass-bottom plate. Displayed is the tracking of mRFP-KDEL-labeled ER tubules using the “Find Fibers” function as average fiber count per cell. Analyzed were 63, 72, 69 cells from shScramble, shRNA2, shRNA3 expressing cells, respectively, from n=3 independent experiments. Two-way ANOVA with Dunnett’s multiple comparison test, **** indicates P < 0.0001).

Although cell growth in YIPF5-KO cells, as well as in YIPF5-KO cells re-expressing either YIPF5 or the YIPF5-I98S mutant, was significantly different from that of MCF10A WT cells after 72 hours (Fig. 1d), the cells remained viable and proliferated. This observation suggests that YIPF5 is not essential for cell viability and is unlikely to play an essential role in the secretory pathway. Loss of the YIPF5-interacting protein IER3IP1 altered ER morphology (Anitei et al., 2024). To investigate whether YIPF5 depletion similarly affected ER morphology, we used a U2OS reporter cell line stably expressing the ER marker RFP-KDEL and shRNAs targeting YIPF5 or a scramble control. Immunoblot analysis confirmed *YIPF5* knockdown, with shRNA3 achieving greater depletion efficiency compared to shRNA2 (Fig. 1f). Using live-cell imaging and high-content analysis, we quantified ER network complexity by measuring the average fiber count (reflecting ER-tubules) per cell as a proxy for network structure. Cells expressing shRNA3 exhibited a significant reduction in ER network complexity compared to control cells (Fig. 1e, g), indicating that loss of YIPF5 disrupts the tubular ER network architecture.

qPCR analysis confirmed increased BiP expression, an established marker of ER stress, in YIPF5-KO cells compared to MCF10A WT cells upon thapsigargin treatment. This increased BIP expression was not rescued by re-expressing YIPF5 WT or I98S (Suppl. Fig. 1c). This is consistent with data from De Franco et al. (De Franco et al., 2020), which demonstrated that YIPF5 deficiency induced ER stress and sensitized β-cells to stress-induced apoptosis. These findings further underscore the critical role of YIPF5 in maintaining ER homeostasis.

Taken together, our results indicate that YIPF5 is present in the early secretory pathway and plays a crucial role in maintaining the integrity of the tubular ER network but is not essential for cell viability or normal growth. MEDS-causing YIPF5 mutants (I98S, L108del, A181V) are not severely mislocalized and do not induce overt morphological changes in ERES or the ERGIC. However, the loss of YIPF5 significantly increases the number of ERES and reduces ER network complexity.

### YIPF5 depletion or mutation alters the secretome, leading to secretion of ER-resident proteins and SURF4 cargoes

Since YIPF5-KO cells are viable and have no obvious defects in ERGIC or Golgi, we hypothesized that YIPF5 might only be involved in ER-export of a subset of cargoes. We therefore determined the secretome and the total proteome of MCF10A WT cells YIPF5 KO-cells or KO cells re-expressing YIPF5 variants using liquid-chromatography-mass spectrometry (LC-MS). To characterize the secretome of these cells, we employed SPECS (Secretome Protein Enrichment with Click Sugars, (Serdaroglu et al., 2017)). This method utilizes the clickable sialic acid analogue ManNAz, which is incorporated into newly synthesized glycoproteins (schematized in Fig. 2a). After LC-MS analysis of supernatants from WT and YIPF5-KO cells, differential abundance analysis was performed using stringent statistical thresholds, leading to the identification of 832 secreted proteins after filtering out nuclear, ribosomal and mitochondrial proteins. (Suppl. Table 1). Proteins whose differential abundance in the secretome mirrored similar changes observed in the corresponding cellular proteome were annotated as “explained by proteome,” indicating that their altered secretion levels were likely reflecting expression changes rather than trafficking effects (Fig. 2b). Of the secreted proteins, 84% were annotated as glycoproteins (https://glycosmos.org/glycoproteins, Suppl. Table 1). Among all detected proteins in YIPF5-KO media, 38,1% remained unchanged, 9.7% reflected changes in total proteome composition, 44,3% showed increased secretion, and 7.8% exhibited reduced secretion compared to WT (Fig. 2b, c; Suppl. Table 1).

**Figure 2:**
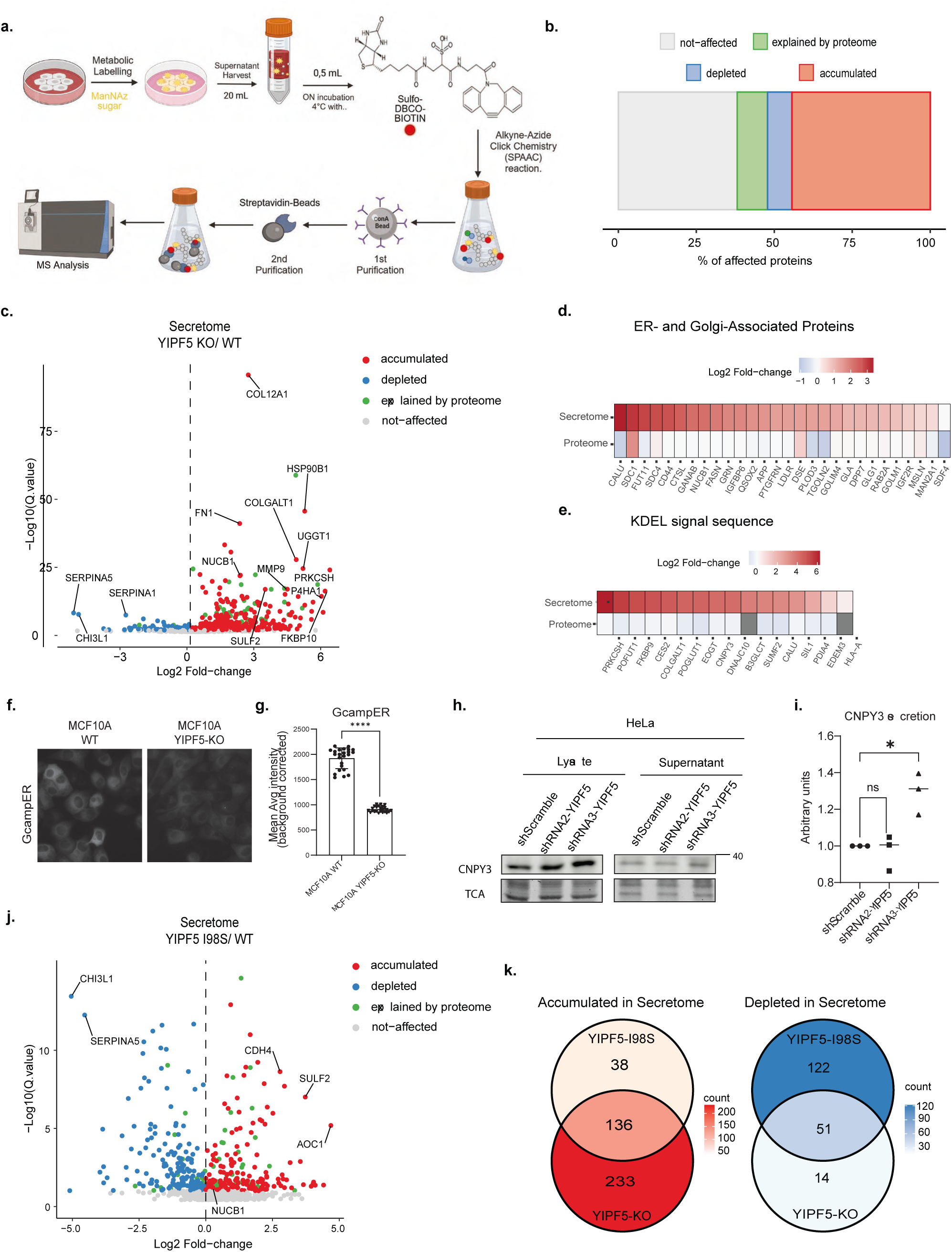
YIPF5 regulates the secretome composition. (a) Schematic representation of the experimental workflow for analyzing the glycoprotein secretome. MCF10A cells were metabolically labeled with the clickable sugar analog ManNAz, followed by a biotin-azide click reaction to tag glycoproteins (n=4). Secreted glycoproteins were concentrated from the culture medium, selectively purified, and identified using mass spectrometry. Image created with BioRender.com. (b) Quantitative comparison of protein abundance in the secretome versus the total proteome of MCF10A cells. Proteins with secretion changes reflecting a similar change in total expression (explained by proteome) (q-value<=0.05) are shown in green. Proteins with increased or decreased secretion independent of total proteome changes are highlighted in red and blue, respectively. Gray (not affected) represents unchanged proteins. (c) Analysis of protein abundance changes in the secretome of YIPF5 KO versus control MCF10A cells (WT). Proteins significantly upregulated (red) or downregulated (blue) in the YIPF5 KO secretome are shown. Proteins with secretion changes explained by total proteome abundance are shown in green, while unaffected proteins are in gray. The vertical axis represents -log10(q-value), and the horizontal axis represents log2(fold change). (d, e) Heatmap depicting proteins showing increased secretion in the YIPF5 KO secretome with a Gene Ontology (GO) annotation for the ER (GO:0007029) and the Golgi apparatus (GO:0005794) (d) or that contain a KDEL-ER retrieval sequence (e). (f) MCF10A WT and YIPF5 KO cells stably expressing the inducible ER-Ca^2+^ sensor GCampER were analyzed using live fluorescence microscopy at identical settings. (g) Mean fluorescence intensity of GCampER was quantified in MCF10A WT and YIPF5 KO cells in 72 wells per cell line from n = 3 independent experiments. Unpaired t-test, two-tailed p < 0.0001. (h) HeLa cells stably expressing shScramble or shRNAs against YIPF5 were transiently transfected with CNPY3-mCherry, supernatants were collected and cell lysates prepared after 24h followed by Western blot analysis using a CNPY3 antibody. A representative blot of n = 3 independent experiments is shown. Source data are available for this figure: SourceData 2. (i) Quantification of CNPY-secretion from (h). Displayed are arbitrary units normalized to the values of shScramble-expressing cells from n = 3 independent experiments. One-way ANOVA with Tukey’s multiple comparisons test, with * indicating p<0.05. (j) Analysis of protein abundance changes in the secretome of YIPF5 KO re-expressing YIPF5 I98S mutant versus control MCF10A cells. Proteins significantly upregulated (red) or downregulated (blue) in the YIPF5 I98S secretome are shown (n=4). Proteins with secretion changes explained by total proteome abundance are denoted in green, while unaffected proteins are in gray. (k) Venn Diagram of protein abundance changes in YIPF5-KO cells and YIPF5-KO re-expressing YIPF5 I98S cells. Proteins with decreased (blue) or increased (red) secretome abundance are shown.

28 proteins enriched in the YIPF5-KO secretome had an ER/Golgi annotation, and 16 carried a KDEL-based ER-retention signal (Fig. 2d). GO-term enrichment analysis of the proteins secreted more in YIPF5-KO cells revealed, among others, significant enrichment in extracellular matrix (ECM) organization, regulation of cell migration, integrin activation, and embryonic development (Suppl. Fig. 2a; Suppl. Table 1). Given the essential role of ECM remodeling and integrin signaling in neuronal migration and cortical development (Franco and Muller, 2013), their dysregulation could contribute to the impaired neuronal positioning observed in microcephaly. Notably, defective ECM composition has been linked to aberrant neurogenesis (Dankovich and Rizzoli, 2022) suggesting that the altered secretome profile upon YIPF5 loss may disrupt the extracellular environment necessary for proper brain development.

Among the KDEL-containing proteins was Calumenin (CALU,Yabe et al., 1997), a known calcium-buffering chaperone (Fig. 2d). This prompted us to investigate whether YIPF5 depletion disrupts ER calcium homeostasis. We identified 66 proteins linked to calcium ion transport (GO:0006816) and calcium ion binding (GO:0005509) that are secreted more in YIPF-KO cells (Suppl. Fig. 2c). To directly assess ER calcium levels, we performed live imaging of MCF10A WT and YIPF5-KO cells stably expressing an inducible version of the genetically encoded calcium sensor GCaMPER (Henderson et al., 2015). YIPF5-KO cells displayed a significant reduction in ER calcium (Fig. 2f, g, expression levels of GCaMPER in Suppl. Fig 2e). Given the central role of ER calcium in protein retention and secretion, its dysregulation may underlie the aberrant secretion of ER-resident proteins observed in YIPF5-deficient cells. Indeed, proteomic profiling identified in addition to Calumenin an increased secretion of ER-resident proteins containing the canonical KDEL retrieval motif, including GDP-fucose protein O-fucosyltransferase 1 (POFUT1, Wang et al., 2001) and CNPY3 (Wakabayashi et al., 2006) (Fig. 2e). This suggests that YIPF5 may regulate KDEL receptor function, similar to its interacting partner IER3IP1 (Anitei et al., 2024). Interestingly, rare CNPY3 variants are associated with early onset epileptic encephalopathy (Ghait et al., 2022; Mutoh et al., 2018), maybe providing a link to the epilepsy phenotype in MEDS2 (De Franco et al., 2020). Among the 369 proteins significantly enriched in YIPF5-KO supernatants, nine proteins including NUCB1, CALU, and SDF4 (CAB45) are well-characterized SURF4 cargoes (Gomez-Navarro et al., 2022; Maldutyte et al., 2025) (for a list of known SURF4-cargoes see Suppl. Table 1 and references therein). Re-expression of WT YIPF5 in YIPF5-KO cells normalized the secretion of some of the dysregulated proteins including NUCB1 (Suppl. Fig. 2d, Suppl. Table 2), indicating that the dysregulation was indeed caused by the loss of YIPF5.

Re-expression of the disease-causing YIPF5-I98S mutant revealed both overlapping and distinct secretion profiles compared to YIPF5-KO cells, suggesting a partial loss-of-function phenotype (Fig. 2k). The volcano plot (Fig. 2j, full list in Suppl. Table 3) provides a quantitative overview of the secretome alterations in YIPF5-I98S mutant cells, revealing a distinct yet overlapping pattern of dysregulated protein secretion compared to YIPF5-KO cells. Notably, 136 secreted proteins, including ECM components and KDEL-containing ER-resident proteins, exhibit increased secretion in both YIPF5-KO and YIPF5 KO re-expressing YIPF5 I98S compared to WT (Fig. 2j, k). However, the degree of dysregulation in YIPF5-I98S cells is less pronounced than in YIPF5-KO cells, supporting a partial loss-of-function phenotype. The Venn diagram further highlights 38 proteins uniquely accumulated and 122 uniquely depleted in the secretome of YIPF5-I98S cells, suggesting an additional gain-of-function phenotype (Fig. 2j). These findings indicate that YIPF5-I98S may still partially engage in cargo selection and retention mechanisms but with reduced efficiency.

The increased secretion of SULF2 in both YIPF5-I98S and YIPF5-KO cells is particularly noteworthy (Suppl. Table 6). Given SULF2’s role in modulating heparan sulfate proteoglycans and its impact on key signaling pathways such as WNT, FGF, and Hedgehog (Rosen and Lemjabbar-Alaoui, 2010), its aberrant secretion may contribute to altered extracellular signaling dynamics relevant for neurodevelopment. In addition, CHI3L1, a secreted glycoprotein involved in neuroinflammation and brain injury response (Jiang et al., 2023), is downregulated in both YIPF5-I98S and YIPF5-KO cells (Suppl. Table 6). Given its role in neuronal survival and repair, CHI3L1 depletion could exacerbate the developmental defects associated with YIPF5 loss.

Next, we validated the secretome analysis by Western Blotting and focused on CNPY3 (Mutoh et al., 2018) as an example of an upregulated protein in the secretome of YIPF5-KO cells (Fig. 2c, e; Suppl. Table 1). HeLa cells stably expressing shRNA against YIPF5 displayed increased secretion of CNPY3 (Fig. 2h, i), as shown by Western Blot, confirming the MS-secretome data.

In summary, our findings provide compelling evidence that YIPF5 depletion or mutation significantly altered the celĺs secretome, causing increased secretion of KDEL-containing proteins and SURF4 cargoes. Furthermore, the impairment of calcium homeostasis and its impact on ER function suggest that YIPF5 is integral to cellular processes that ensure proper protein sorting and secretion. These changes in the secretome are associated with pathways relevant to neuronal migration and brain development, implicating YIPF5 as a key player in neurodevelopmental processes, with potential contributions to disorders such as microcephaly. Notably, the partial loss-of-function phenotype observed in YIPF5-I98S mutant cells suggests a pathomechanism for MEDS2.

### YIPF5 and SURF4 interact

The enhanced secretion of potential SURF4 cargoes in the absence of YIPF5 prompted us to examine the physical interaction between the two proteins. Co-immunoprecipitation assays demonstrated that GFP-tagged YIPF5 successfully pulled down FLAG-tagged SURF4, while GFP-SURF4 co-precipitated endogenous YIPF5 (Fig. 3a, Suppl. Fig. 3), confirming their association. High-resolution Airyscan microscopy revealed that GFP-SURF4 and YIPF5 co-localized within the secretory pathway (Fig. 3b). YIPF5 depletion caused a pronounced redistribution of SURF4 from the ER to post-ER compartments in MCF10A YIPF5-KO cells stably expressing GFP-SURF4 (Fig. 3c, Video 1). Immunostaining with the ERES marker SEC31A (Fig. 3d) revealed a reduced co-localization between SURF4 and ERES in YIPF5-KO cells (Fig. 3e). Interestingly, the deletion of YIPF5 induced the formation of SURF4-positive, SEC31A-negative tubules emerging from ERES (Fig. 3d). Fig. 3f, g display increased GFP-SURF4 colocalization with the cis-Golgi marker GM130 in YIPF5-KO cells.

**Figure 3.**
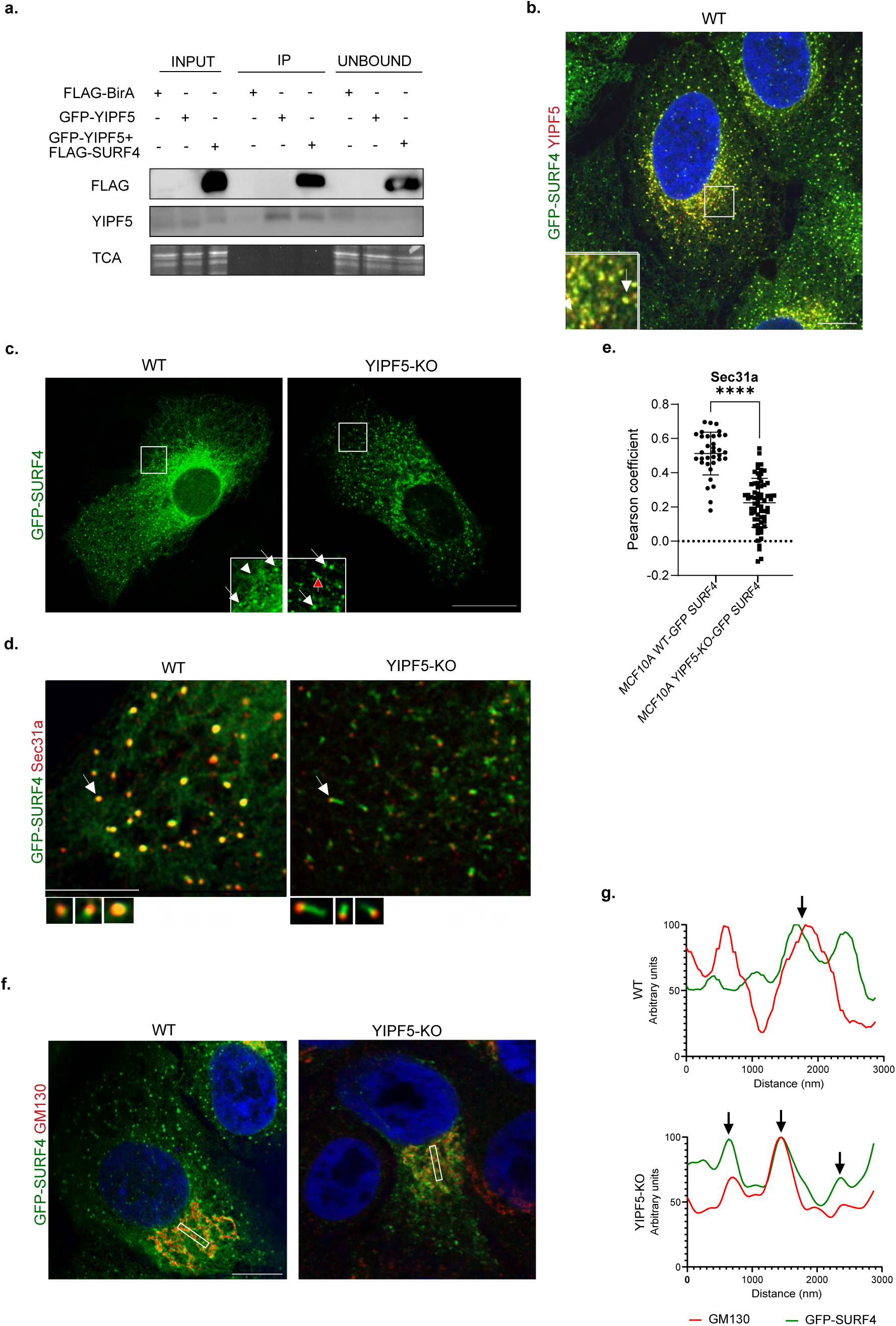
Deletion of YIPF5 induces relocalization of SURF4. (a) HeLa cells were transiently transfected with plasmids expressing GFP-YIPF5 and FLAG-SURF4 or FLAG-BirA (negative control) and cell lysates prepared after 24h. Lysates were immunoprecipitated using GFP-TRAP beads, followed by Western blot analysis with indicated antibodies. Trichloroacetic acid (TCA) precipitation was used as a loading control to ensure equal protein amounts across samples. A representative blot of n = 3 independent experiments is shown. Source data are available for this figure: SourceData F3. (b) MCF10A WT cells stably expressing GFP-SURF4 (green) were fixed, stained for YIPF5 (red) and imaged by Airyscan super-resolution microscopy. A Z-projection is shown. The insert shows a magnification of the boxed area. Arrows indicate co-localization of YIPF5 and SURF4. Nuclei were stained with Hoechst (blue). Scale bar: 10 µm. (c) MCF10A WT or YIPF5 KO cells stably expressing GFP-SURF4 (green) were fixed and analyzed by Airyscan super-resolution microscopy. Insets show magnified views of boxed areas. The arrowhead depicts localization in the tubular ER-network, the white arrows localization to ERES and the red arrow to post-ERES tubular structures. Scale bar 10 µm. (d) MCF10A WT or YIPF5 KO cells stably expressing GFP-SURF4 (green) were fixed, stained with the ERES marker SEC31A (red) and analyzed by Airyscan super-resolution microscopy. Magnified views of typical ERES (arrows) are shown below. Scale bar: 2 µm. (e) Quantification of the co-localization between SEC31A and GFP-SURF4 in cells from (d). Displayed is the mean Pearson coefficient ± SEM from n = 3 independent experiments and 34 WT and 66 YIPF5-KO cells. Unpaired t test, **** indicates p < 0.0001). (f) MCF10A WT or YIPF5 KO cells stably expressing GFP-SURF4 (green) were fixed and co-stained with Golgi marker GM130 (red). Images were acquired using Airyscan super-resolution microscopy. Scale bar: 10 µm. (g) Fluorescence intensity blots indicating the co-localization of GFP-SURF4 and GM130 in the selected regions of interest for each condition.

### YIPF5 depletion induces SURF4-positive tubules and promotes CAB45 secretion

To further characterize the SURF4-GFP-positive tubules, we performed live-cell imaging in MCF10A YIPF5-KO cells expressing GFP-SURF4 and Sec24C-mCherry. Long, dynamic SURF4-positive tubules originated from ERES, often extending over several μm (Fig. 4a). A time-lapse video (Video 2), captured using super-resolution microscopy, clearly demonstrates the extension of these tubules from ERES over time (Fig. 4a). Fixed-cell imaging further revealed that the GFP-SURF4-positive tubules are ERGIC-53 positive (Fig. 4b), also visualized by the intensity profile plots of two regions of interest (ROIs) (Fig. 4c). In addition, the GFP-SURF4 tubules are also positive for RAB1b, a small GTPase known to regulate ER-to-Golgi trafficking and vesicle tethering at the ERGIC (Peter et al., 1994; Westrate et al., 2020) (Suppl Fig. 3e and Video 3). Given the observed alterations in GFP-SURF4 trafficking in the absence of YIPF5, we investigated the functional consequences on the secretion of CAB45 a well-established SURF4 cargo (Gomez-Navarro et al., 2022; Maldutyte et al., 2025) in HeLa cells stably expressing shYIPF5 or scramble shRNA. We made use of the sensitivity of N-glycosylated proteins to Endoglycosidase H (EndoH) to assess the transport of Rush-CAB45-GFP. Sensitivity (sen) is indicative of a localization in pre-Golgi compartments whereas EndoH resistance (res) is indicative for a Golgi/post-Golgi localization. Quantification of the ratio EndoH res/sen is an indicator of transport through the secretory pathway and revealed a significant increase at 20 and 30 minutes in YIPF5-depleted cells compared to controls (Fig. 4e). Supporting these data, more Rush-CAB45-GFP accumulated in the supernatant of in YIPF5-depleted cells (Fig. 4f). This suggests an enhanced ER-to-Golgi trafficking of Rush-CAB45-GFP via ERES-derived SURF4-tubules in YIPF5-depleted cells, in line with an increase of Rush-CAB45-GFP in the secretome. Collectively, these findings establish YIPF5 as a critical determinant of SURF4 localization and as a negative regulator of SURF4-dependent ER-Golgi transport.

**Figure 4.**
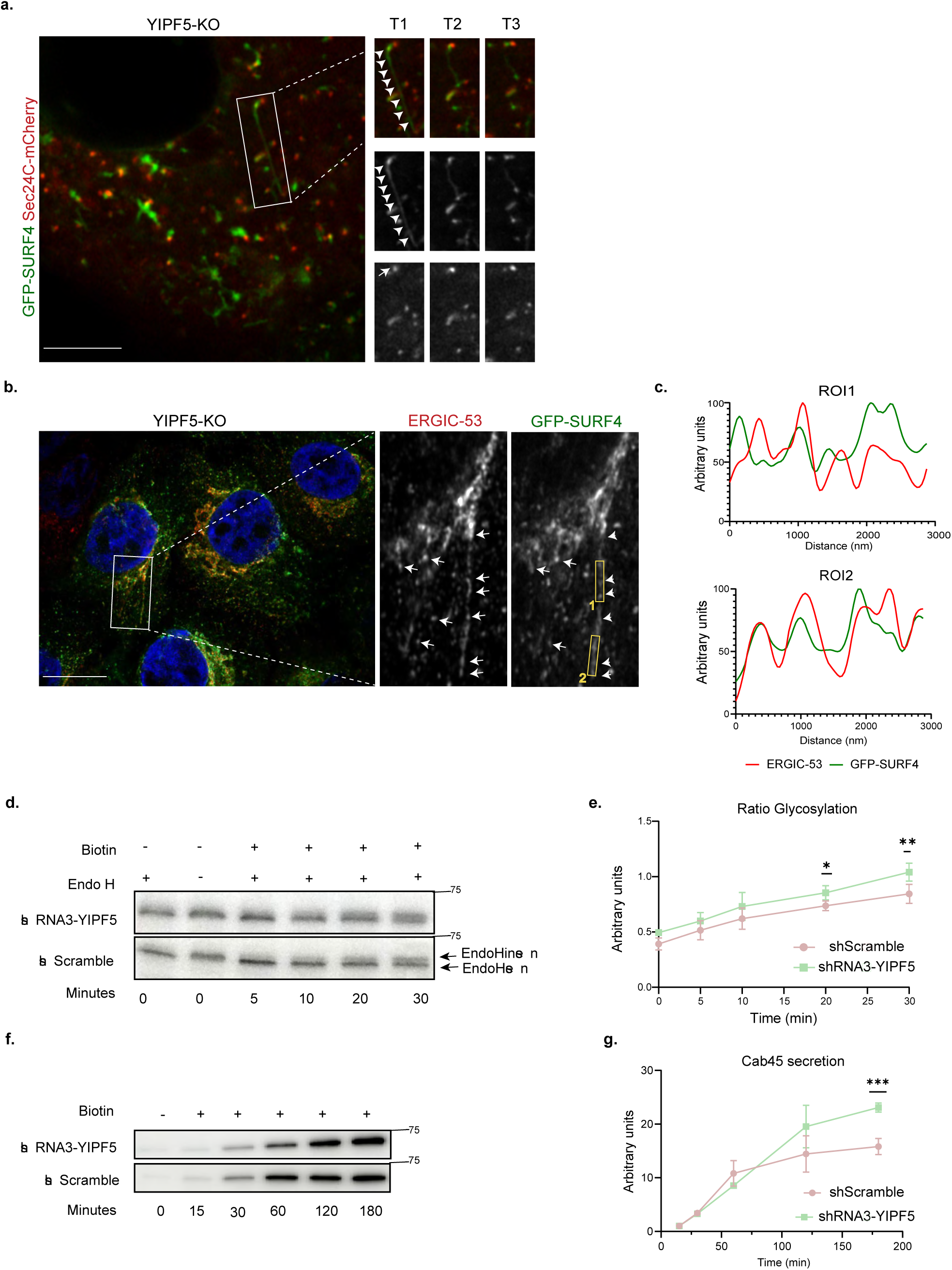
YIPF5 depletion induces the formation of dynamic SURF4-positive tubules and accelerates the secretion of the SURF4 cargo CAB45. (a) Live-cell Airyscan super-resolution microscopy of MCF10A YIPF5-KO cells stably expressing GFP-SURF4 (green) and transfected with SEC24C-mCherry (red), marking ERES. Right: Magnifications of the boxed area at three different timepoints (T1-3, 0,5 seconds interval). Arrowheads depict a tubular structure emanating from an ERES (arrow). Arrowheads indicate the dynamic extension of GFP-SURF4-positive tubules, while the full arrow point to a Sec24c-labeled ERES. Scale bar: 2 µm. (b) Airyscan imaging of fixed YIPF5-KO cells expressing GFP-SURF4 (green) and immunostained for ERGIC-53 (red). Enlarged regions show ERGIC-53-positive labeling (arrows) along SURF4-positive tubules (arrowheads). Scale bar: 10 µm. (c) Intensity profile plots of two regions of interest (ROI1, 2) from (b). (d, f) HeLa cells transfected with RUSH-CAB45-GFP and stably expressing either shScramble or shRNA3-YIPF were treated with Biotin (40 µg/ml) for indicated timepoints. (d) Cell lysates were treated with Endoglycosidase H (Endo H) before analysis by Western Blot using anti-GFP antibody. Arrows indicate EndoH resistant (res) or sensitive (sen) forms of RUSH-CAB45-GFP. (f) Supernatants were subjected to GFP-trap pull-down, separated by SDS-PAGE, blotted and probed with anti-GFP antibody. (e) Quantification of (d). Displayed is the ratio EndoH_res_/EndoH_sen_ as mean ± SD of n=3 experiments. Statistical analysis: multiple paired t-test with two-stage step-up correction (Benjamini, Krieger, and Yekutieli). *p ≤ 0.05, **p ≤ 0.01. (g) Quantification of (f). Displayed is the mean secreted GFP-CAB45 as arbitrary units normalized to 15 minutes, ± SEM of n=4 experiments. Statistical test: multiple unpaired t test with Welch correction. Source data are available for this figure: SourceData F4.

### YIPF5 regulates surface trafficking of membrane proteins

After establishing the role of YIPF5 in the trafficking of soluble proteins, we investigated its function in the transport of membrane proteins. To this end, we performed cell surface biotinylation followed by streptavidin pull-down and LC-MS analysis to profile the surfaceome of MCF10A wild-type (WT) cells, YIPF5 knockout (KO) cells, and KO cells re-expressing various YIPF5 constructs (outlined in Fig. 5a). In parallel, the total proteome was analyzed by LC-MS and compared with the secretome.

**Figure 5:**
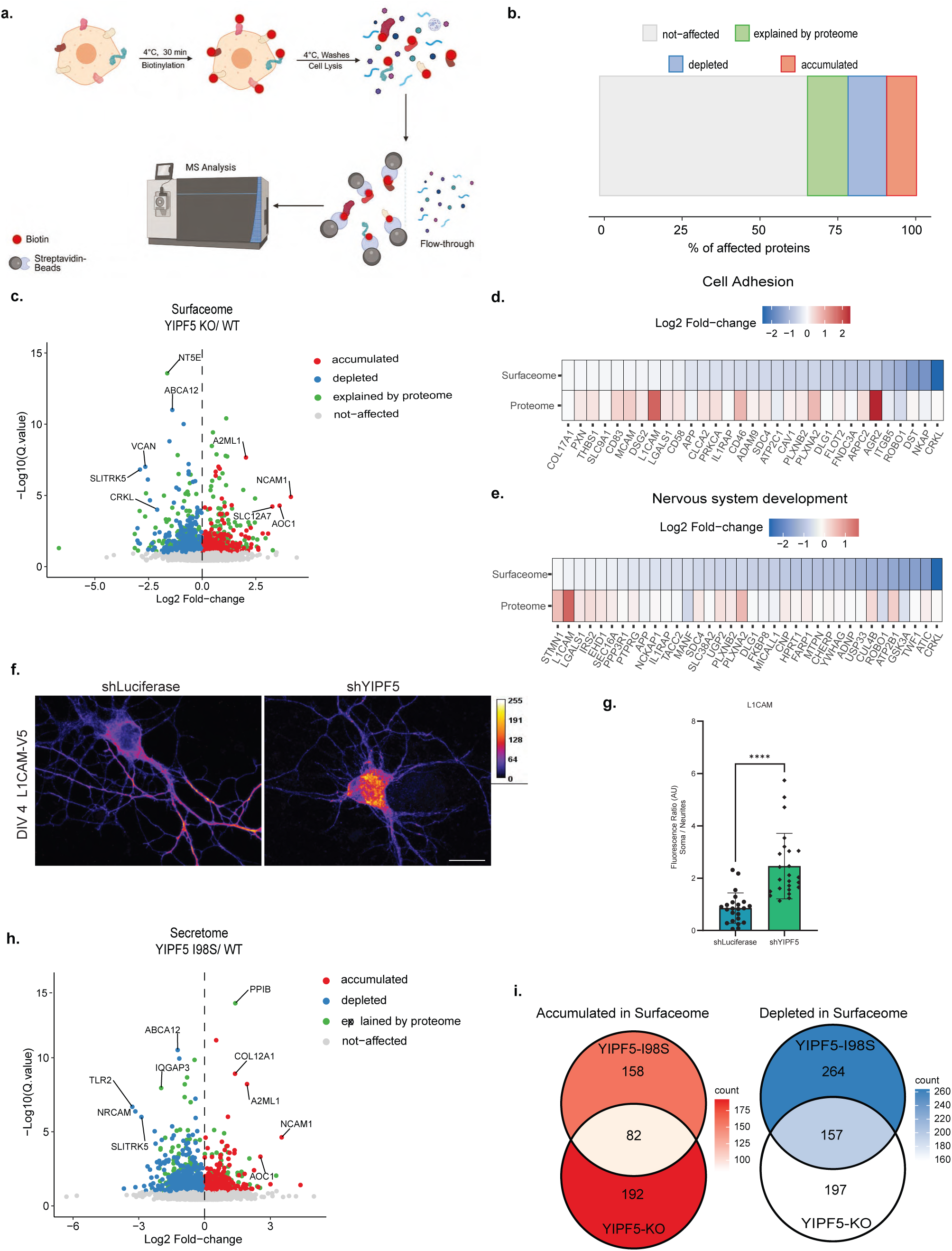
YIPF5 regulates cell surface proteome dynamics and impacts neuronal protein trafficking. (a) Schematic of surface biotinylation workflow in MCF10A cells. MCF10A cells were labeled with biotin to capture surface proteins, followed by streptavidin-based purification and mass spectrometry analysis (n=4). Image created with BioRender.com. (b) Quantitative comparison of protein abundance changes on the cell surface relative to the total proteome in MCF10A YIPF5-KO cells vs. WT cells. Proteins with increased surface expression are highlighted in red (q-value<0.1), while those with decreased surface expression are shown in blue (q-value<0.1). Proteins whose surface changes are explained by total proteome abundance are denoted in green, and unaffected proteins are in gray. (c) Analysis of protein abundance changes on the cell surface of YIPF5 KO versus control MCF10A WT cells. Proteins significantly upregulated (red, q-value<0.1), or downregulated (blue, q-value<0.1) in YIPF5 KO cells are shown. Proteins with surface expression changes explained by total proteome abundance are denoted in green, while unaffected proteins are in gray. The vertical axis represents -log10(q-value), and the horizontal axis represents log2fold change. (d, e) Heatmap displaying cell adhesion-related proteins (GO:0007155) significantly reduced on the surface of YIPF5 KO MCF10A cells compared to WT (d) or associated with nervous system development (GO:0007399) with reduced surface expression in YIPF5 KO cells. Proteins with decreased surface abundance are shown in varying shades of blue, total proteome abundances in red, with darker tones indicating greater depletion. (f) DIV4 rat hippocampal neurons were co-transfected with L1CAM-V5 and either shLuciferase (control) or shYIPF5, cultured with biotin for 48h, fixed and stained with anti-V5 antibody and imaged by confocal microscopy. Representative confocal images are shown. Fluorescence intensity is displayed using a Turbo heatmap (dark blue: zero intensity; yellow/red: maximum intensity). (g) Quantification of L1CAM fluorescence intensity in soma versus neurites (n = 24 neurons for shYIPF5, n = 23 neurons for shLuciferase; n = 4 independent experiments). Statistical significance was determined using an unpaired t-test with Welch’s correction (**** indicates p < 0.0001). (h) Analysis of protein abundance changes on the cell surface of YIPF5 I98S mutant versus WT MCF10A cells. Proteins significantly upregulated (red, (q-value<0.1) or downregulated (blue (q-value<0.1) in the YIPF5 I98S mutant are shown. Proteins with surface expression changes explained by total proteome abundance are denoted in green, while unaffected proteins are in gray. (i) Overlap and exclusivity in protein abundance changes on the cell surface of MCF10A YIPF5 I98S mutant cells compared to YIPF5 KO cells. Proteins with decreased (blue) or increased (red) surface presence are shown. The intersection highlights proteins with consistent changes in both YIPF5 I98S and KO cells.

A total of 2,902 proteins were identified at the cell surface. After excluding nuclear, ribosomal and mitochondrial proteins, 630 biotinylated proteins showed significant alterations in surface abundance in YIPF5-KO cells relative to WT controls. Similarly to the secretome, proteins were annotated as “explained by proteome” when their direction of change at the cell surface matched that observed in the total proteome, indicating a consistent regulation at both levels. (Fig. 5b, c; Suppl. Table 4). Among all proteins detected on the surface of YIPF5-KO cells, 65.6% remained unchanged, 12.78% reflected changes in total proteome composition, 9.44% showed increased accumulation at the plasma membrane, and 12.19% exhibited reduced surface localization compared to WT (Fig. 5b; suppl. Table 4.)

Gene Ontology (GO) annotation of surface-depleted proteins in YIPF5-KO cells revealed significant enrichment in terms related to cell adhesion, system development, neuron differentiation, and neuron projection development, with cell adhesion emerging as the most significantly affected category (suppl. Fig. 5; Suppl. Table 4). Heatmap analysis further highlighted the depletion of proteins associated with cell adhesion and nervous system development in YIPF5-KO cells compared to WT controls (Fig. 5d, e). Among these, key neuronal cell adhesion molecules, including L1CAM (Linneberg et al., 2019) and Plexin-A2 (PLXNA2) (Zhao et al., 2018), exhibited increased abundance in cell lysates but were significantly reduced at the plasma membrane in YIPF5-KO cells, indicating a defect in their transport to the cell surface. To validate these findings, we expressed a RUSH(retention using selective hooks, (Boncompain et al., 2012))-based L1CAM-V5 construct in primary rat hippocampal neurons transfected with either a control (shLuciferase) or an shRNA vector targeting YIPF5 and incubated with biotin. In shLuciferase-expressing neurons, L1CAM localized predominantly to neuritic surfaces, whereas in shYIPF5-transfected neurons, L1CAM accumulated in the soma in an ER-like pattern, indicative of a trafficking defect. Quantification confirmed an increased soma-to-neurite ratio of L1CAM staining in shYIPF5-transfected neurons, suggesting that YIPF5 is required for efficient transport of L1CAM to neuritic surfaces (Fig. 5f, g). KD-efficiency is shown in suppl. Fig. 4b.

Similar defects in surface trafficking were observed in YIPF5-KO cells re-expressing the disease-causing YIPF5-I98S mutant, mirroring the changes detected in the YIPF5-KO surfaceome (Fig. 5h; suppl. Table 5). Among the 158 upregulated surface proteins in YIPF5-I98S cells, 82 overlapped with YIPF5-KO cells, while among the 264 downregulated proteins, 157 were shared with YIPF5-KO cells (Fig. 4i; suppl. Table 5). Notably, CRKL (Matsuki et al., 2008), SLITRK5 (Liu et al., 2022), and ROBO1 (Yeh et al., 2014), all crucial for neuronal development, were depleted from the surface in both YIPF5-KO and YIPF5-I98S cells. Given their established roles in axon guidance and cortical development, their mislocalization may contribute to the microcephaly phenotype associated with YIPF5 mutations. Interestingly, PlexinD1, a protein involved in neuronal migration termination (Sawada et al., 2018), was successfully trafficked to the surface in YIPF5-I98S-expressing cells but not in YIPF5-KO cells (Suppl. Table 7), indicating that the I98S mutant retained partial functionality for selected substrates. These findings reveal that YIPF5 plays a critical role in membrane protein trafficking, particularly for neuronal adhesion and axon guidance molecules. The inability to efficiently transport these proteins to the plasma membrane in YIPF5-KO and YIPF5-I98S cells suggests a failure in intracellular trafficking, which may underline the neurodevelopmental defects associated with YIPF5 mutations.

### YIPF5 deficiency impairs cell migration *in vitro* and neuronal migration *in vivo*

Having found that loss or mutation of YIPF5 alters the homeostasis of the surfaceome, particularly affecting proteins linked to migration, neurodevelopment, and cellular adhesion, we sought to determine whether YIPF5 played a role in cellular migration *in vitro*. To this end, we performed a wound healing assay using live cell imaging with the IncuCyte system (Fig. 6a). WT MCF10A cells closed the scratched area by only 20% after 15 h, whereas YIPF5-KO cells or YIPF5-KO cells re-expressing the pathogenic I98S YIPF5 mutant exhibited a significantly increased wound closure, reaching 50% after 15 h. Similarly, YIPF5-KO cells re-expressing YIPF5 at either low or high levels displayed enhanced migration, suggesting that both deletion and overexpression of YIPF5 accelerate wound healing. In line with these findings, stable knockdown of YIPF5 in U2OS cells also promoted wound closure, particularly in cells expressing the more efficient shRNA3, further supporting the role of YIPF5 in cellular migration (Suppl Fig. 6).

**Figure 6.**
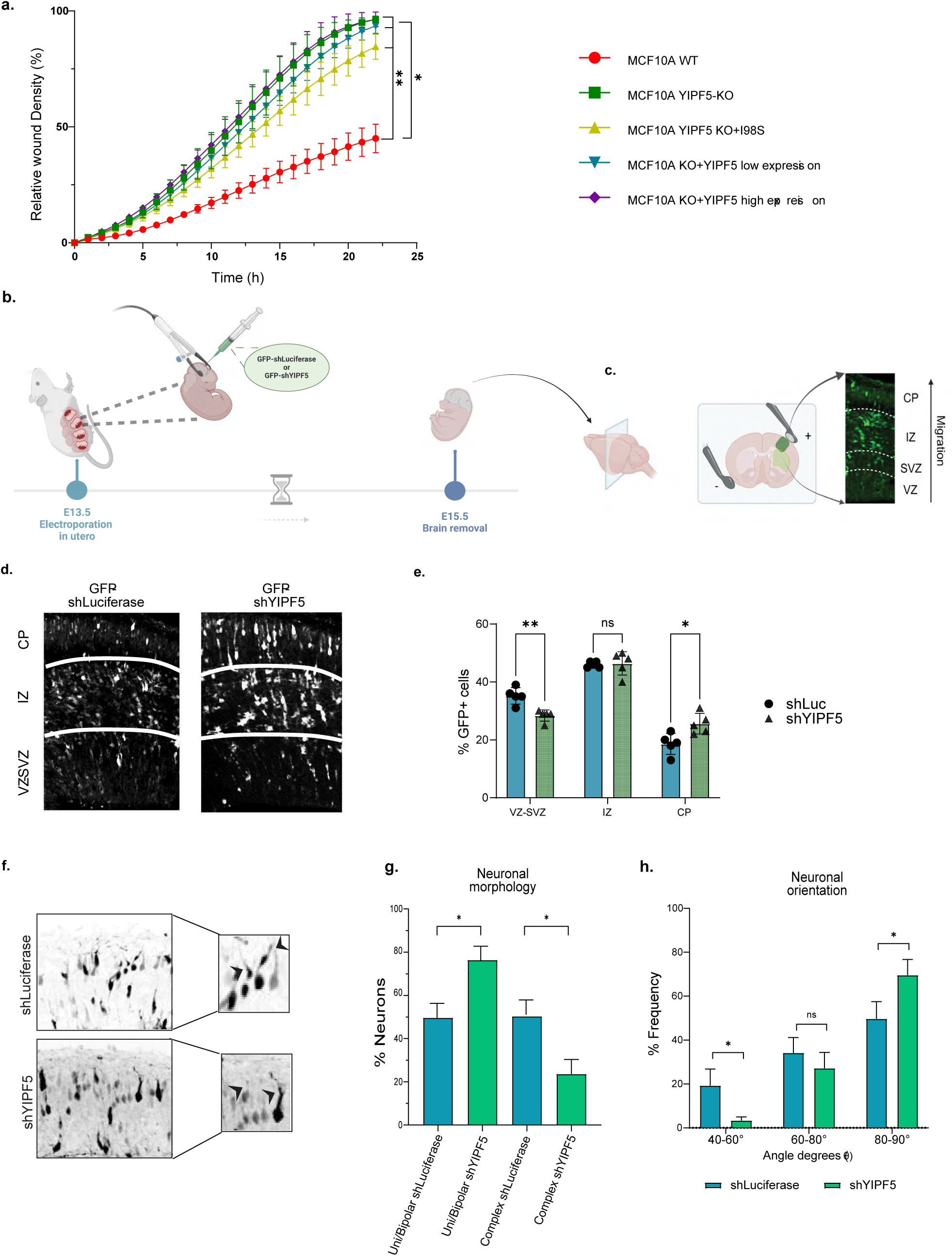
YIPF5 regulates cellular and neuronal migration in vitro and in vivo. (a) MCF10A WT, YIPF5-KO, and YIPF5-KO cells re-expressing either the I98S YIPF5 mutant or WT YIPF5 at low or high levels were plated in Incucyte® Imagelock 96-well plates and a wound healing assay performed using the Incucyte® Wound Maker 96-Tool on the IncuCyte system. Displayed is the relative wound density at different time points as mean values ± SEM from three biological replicates. Statistical significance was evaluated using a two-way ANOVA with Tukey’s multiple comparisons test, with ** denoting p<0.01 and * indicating p<0.05. (b) Schematic representation illustrating the in-utero electroporation procedure at embryonic day 13.5 (E13.5) with a GFP-labeled plasmid encoding either shLuciferase or shYIPF5, followed by brain removal at E15.5. (c) Scheme of a coronal section of a mouse embryonic brain schematizing the position of the electrodes, the ventricles and the developing neocortex with the classification into distinct zones: ventricular zone (VZ), subventricular zone (SVZ), intermediate zone (IZ), and cortical plate (CP). This classification traces the cortical migration pattern of GFP+ electroporated cells. Illustrations created with BioRender.com using an IHC image deriving from our murine brain. (d) Murine brains electroporated at E13.5 with plasmids expressing control shLuciferase or shYIPF5 were harvested at E15.5, fixed, sectioned with a vibratome, and immunostained for GFP to visualize transfected cells. Nuclear staining with Hoechst (not shown) facilitated the differentiation of brain regions, including VZ, SVZ, IZ and CP. Representative images are shown. (e) Quantification of the proportion of EGFP+ cells within specified cortical regions. Displayed are the means ± SEM (shLuciferase, n = 5 brains, 225 EGFP+ cells; shYIPF5, n = 5 brains, 257 EGFP+ cells). Statistical analysis: multiple unpaired t test. (f) Morphometric analysis of CP-localized neurons of shLuciferase and shYIPF5 electroporated brains. Images are maximum projections of sequential Z-sections, the inverted EGFP signal is depicted in black. (g) quantification of neuron morphologies from shLuciferase (n=6 brains, 6 sections, 93 EGFP+ cells) and shYIPF5 (n=5 brains, 6 sections, 71 cells) groups. Neurons were categorized into two groups: unbranched uni/bipolar and exhibiting complex arborizations. Mean values with standard deviation are shown, statistical analysis: one-way ANOVA with Šídák’s multiple comparisons test, *p <0.05. (h) Leading process orientation analysis of CP-localized neurons from brain sections as shown in f. Arrowheads display examples of tilted neurites. The angles between the leading neurites of EGFP+ neurons and the CP surface were calculated using the measurement feature in ImageJ. The frequency of angle values was categorized into specified ranges, expressed as percentages of the total number of values. The sample sizes comprised 92 EGFP+ cells from 6 sections from 6 brains for the shLuciferase group and 96 EGFP+ cells from 9 sections from 5 brains for the shYIPF5 group. Independent unpaired t-tests were conducted for the three angle ranges, with a significance level set at *p < 0.05.

To explore the function of YIPF5 in neuronal migration *in vivo*, we performed *in utero* electroporation at E13.5 using EGFP-expressing plasmids carrying either control shRNA or shRNAs targeting YIPF5 into neural progenitors of the lateral cortex (Fig. 6b-c). Immunoblotting confirmed efficient KD of YIPF5 achieved by the shRNA in NIH3T3 cells (Suppl. Fig. 5f). Quantification of EGFP⁺ neurons in coronal brain sections of E15.5 brains revealed a significant redistribution of cells across cortical layers: YIPF5 depletion led to a reduction of electroporated, EGFP^+^ cells in the ventricular and subventricular zones (VZ-SVZ) and a concomitant increase in the cortical plate (CP), consistent with increased migration of newborn neurons (Fig. 6e, f). Furthermore, Morphometric analysis of neurons localized to the cortical plate (CP) revealed a significant increase in unbranched bipolar cells, accompanied by a decrease in neurons with complex arborizations (Fig. 6g, h). This change indicates impaired morphological maturation and suggests that YIPF5 could play a crucial role in the transition between multipolar and bipolar neuronal stages during cortical migration (Kon et al., 2017). Additionally, an analysis of leading process orientation in the same area showed a decrease in the variability of neurite angles, with a notable concentration of neurons oriented at 80°–90° relative to the CP in the case of shYIPF5 (see Fig. 6i). Collectively, these findings highlight a critical role for YIPF5 in orchestrating proper neuronal positioning and morphology, possibly through its regulation of the secretory pathway and surface protein dynamics.

## Discussion

We here demonstrate that the small polytopic ER-membrane protein YIPF5 is involved in ER-exit of a subset of secreted and PM proteins. Interestingly, the absence of YIPF5 does not generally increase or decrease surface transport or secretion. While ER-resident proteins with a C-terminal KDEL retention signal and cargoes of the ER-exit receptor SURF4 were generally secreted more, many membrane proteins involved in cellular migration and neuronal development were transported less to the surface. In the developing brain this has drastic consequences, indicated by the microcephaly in the fatal MEDS2 syndrome caused by loss-of-function mutations in YIPF5 (De Franco et al., 2020). How loss or mutation of YIPF5 results in selective increase of transport of some proteins and decreased transport of other proteins is not yet fully understood but could result from a mixture of direct and indirect effects. The increased ER-export of SURF4 cargoes is probably a direct effect, since YIPF5 colocalized with SURF4 in superresolution microscopy and both directly interacted in co-immunoprecipitation experiments. Also, GFP-SURF4 was shifted from a more ER-ERES localization to post-ERES localization in YIPF5-KO cells. Since loss of YIPF5 increased secretion of SURF4-cargoes, we propose that it functions as a negative regulator of SURF4-dependent ER-export, maybe as part of a quality control system. Abolition of this negative regulation might lead to loss of ER-retention and retrieval mechanisms, resulting in aberrant secretion of normally ER-resident proteins, including those that are involved in Ca^2+^-homeostasis. Loss of Ca^2+^ in the ER was shown previously to result in secretion of KDEL-containing ER-resident proteins (Booth and Koch, 1989; Trychta et al., 2018). This might in turn result in dysregulation of Ca^2+^-dependent chaperoning in the ER resulting in the accumulation of unfolded or misfolded proteins, eventually leading to the decreased ER-export of some membrane proteins. YIPF5 could not only negatively regulate SURF4-dependent ER-export but also other ER-export pathways, explaining the increased secretion of many non-SURF4 cargoes. Alternatively, loss or mutation of YIPF5 could primarily cause a reduction in ER-Ca^2+^-levels, resulting in release of KDEL-containing ER-proteins and dysregulation of ER-export. SURF4 was directly implicated in Ca^2+^-homeostasis via the store operated calcium channel SOCE, although ER-Ca^2+^ was not measured (Fujii et al., 2012; Picon-Pages et al., 2023). SURF4 transports proinsulin out of the ER (Saegusa et al., 2022), but why loss of function YIPF5-mutations in MEDS2 cause impaired not increased insulin secretion (De Franco et al., 2020) needs to be further elucidated.

In the absence of YIPF5, SURF4 exited the ER in long thin tubules and secretion of Cab45, a SURF4-cargo (Maldutyte et al., 2025), was increased. A possible molecular mechanism of this enhanced ER-exit could be the fact that YIPF5 interacts via its N-terminus with COPII components (Ran et al., 2019; Tang et al., 2001) and might control COPII activity and/or specificity. Alternatively, it might control the expansion of the transport carrier, allowing more time for selection and control of export cargo. Loss or mutation of YIPF5 might increase COPII activity at ERES (activity here meaning on/off rates, shown to be related to export activity (Forster et al., 2006). Concomitantly, transport carrier formation would be enhanced, resulting in the formation of very long carriers that eventually pinch off towards the Golgi. In future work this model could be tested by FRAP-experiments at ERES to determine COPII-dynamics. Instead, YIPF5 could be involved in retrograde transport, possibly COPI-dependent, from emerging ER-Golgi transport carriers, to recycle membrane and ER-exit machinery back to the ER. Depletion of YIPF5 could cause a defect in this mechanism, resulting in the long tubules that we observed.

SURF4 was shown before to exit ERES in long, thin tubules, termed tERGIC, that are ERGIC53 negative and RAB1b positive (Yan et al., 2022). How these tubules are related to the SURF4-tubules observed here remains an open question. We didńt observe extended GFP-SURF4 tubules in WT cells, only in YIPF5-deficient MCF10A or Hela cells. Also, our SURF4-tubules are ERGIC53 and RAB1b positive, unlike the ERGIC53 negative and RAB1A/B positive tERGIC (Yan et al., 2022), suggesting they are different structures.

GFP-SURF4 tubules emerged from COPII-labelled ERES and, after scission, moved towards the Golgi, suggesting they are bona-fide, albeit extended ER-export transport carriers. COPII always stayed at ERES, supporting the recently proposed model by us and others that the role of COPII is more of a collar around the neck connecting ER and ERES than an actual vesicular coat (Shomron et al., 2021; Weigel et al., 2021). Our findings raise the possibility that the secretome of YIPF5-KO cells may contain yet-undiscovered SURF4 cargoes and suggests a functional interplay between YIPF5 and SURF4, potentially implicating YIPF5 in cargo selection and ER export mechanisms.

Possible consequences of the enhanced secretion of ER-resident proteins could be two-fold. A disbalance of chaperones in the ER could have effects on protein folding, compromising for example ER-quality control and consequently ER-export of misfolded proteins with detrimental consequences at their final destination. Or, lack of ER-chaperones due to unwanted secretion could affect folding capabilities in the ER, resulting in accumulation of unfolded proteins. This might explain the reduced transport of selected proteins to the cell surface. In the future it will be interesting to analyze if those proteins we found downregulated on the PM are the ones with the highest need of chaperone support, maybe because of many transmembrane domains or essential assembly in sub-complexes prior to ER-export. One the other hand, the secreted ER-resident proteins could have functions at the extracellular space, for example HSP90B1 which is more secreted in YIPF5-depleted cells and has a role in neurodevelopment (Miller and Fort, 2018).

The increased migration we observed *in-vitro* in cultured YIPF5 KD cells and *in-vivo* in mouse embryos suggests that faster and/or overmigration of neuronal precursor cells and/or newborn neurons is a novel cause of microcephaly. Microcephaly is mostly attributed to reduced proliferation of neuronal stem cells/progenitors or to increased cell death of neuronal precursors (Heide and Huttner, 2023; Jean et al., 2020), but also to defects in migration and morphology of neurons (Asselin et al., 2020; Ruiz-Reig et al., 2022). Future work, ideally using a mouse model for MEDS2 will shed light on the precise mechanism. Variants in IER3IP1 cause MEDS1 (Abdel-Salam et al., 2012; Poulton et al., 2011; Shalev et al., 2014). IER3IP1 interacts with YIPF5 and we showed that KO or mutation of IER3IP1 causes changes in secretome and surfaceome similar to what we observed here for YIPF5 (Anitei et al., 2024). Similarly, in the absence of IER3IP1, proteins important for neuronal development were less abundant at the surface and KDEL-containing proteins were increasingly secreted, suggesting very similar pathomechanisms for MEDS1 and 2.

## Materials and methods

*For Antibodies and primers see supplementary table 8*.

### Plasmids

| Name | Donor/origin | Reference |
| --- | --- | --- |
| **LeGO-iG2** | Addgene #27341 was a gift from Boris Fehse (http://n2t.net/addgene:27341 ; RRID:Addgene_27341) | (Weber et al., 2008) |
| **Rab1b-mCherry** |  | (Shomron et al., 2021) |
| **mCherry-C1-Sec24C** |  | (Shomron et al., 2021) |
| **pFLAG-SURF4** | Addgene #170680 was a gift from Ke Xu (http://n2t.net/addgene:170680 ; RRID:Addgene_170680) | (Yan et al., 2022) |
| **SF-LV-shLuciferase-mirE** | Kind gift from Lenhard Rudolph | (Wang et al., 2012) |
| **pplss-mRFP-KDEL** | Kind gift from Eric Snapp | (Snapp et al., 2006) |
| **FUGW_Strep-**<br>**li_L1CAM-APEX2-**<br>**V5-SBP** | This paper |  |
| **psPAX2** | Addgene, #12260 was a gift from Didier Trono (http://n2t.net/addgene:12260 ; RRID:Addgene_12260) |  |
| **pMD2.G** | Addgene, #12259 was a gift from Didier Trono (http://n2t.net/addgene:12259 ; RRID:Addgene_12259) |  |
| **pMOS003-lenti-**<br>**CMV-GCaMPer** | was a gift from Adam Cohen & Brandon Harvey (Addgene plasmid # 65227 ; http://n2t.net/addgene:65227 ; RRID:Addgene_65227) | (Henderson et al., 2015) |
| **pInd20-GCampER** | This paper |  |
| **LeGO_iG2_EGFP_**<br>**YIPF5WT** |  | (Cramer et al., 2023) |
| **LeGO_iG2_EGFP_**<br>**YIPF5-A181V** | This paper |  |
| **LeGO_iG2_EGFP_**<br>**YIPF5-I98S** | This paper |  |
| **LeGO_iG2_EGFP_**<br>**YIPF5-delL106** | This paper |  |
| **LeGO_iG2_EGFP_**<br>**YIPF5-W218R** | This paper |  |
| **pcDNA3.1_FLAG-**<br>**hYIPF5wt** | This paper |  |
| **pcDNA3.1_V5-**<br>**hYIPF5wt** | This paper |  |
| **SGEP** | Kind gift from Johannes Zuber | (Fellmann et al., 2013) |
| **SGEP_shRNA1-**<br>**YIPF5** | This paper |  |
| **SGEP_shRNA2-**<br>**YIPF5** | This paper |  |
| **SGEP_shRNA3-**<br>**YIPF5** | This paper |  |
| **SF-LV-shYIPF5-**<br>**mirE_mouse** | This paper |  |
| **SF-LV-shYIPF5-2-**<br>**mirE_mouse** | This paper |  |
| **SF-LV-shYIPF5-3-**<br>**mirE_mouse** | This paper |  |
| **SF-LV-shYIPF5_1-**<br>**mirE_rat** | This paper |  |
| **SF-LV-shYIPF5_2-**<br>**mirE_rat** | This paper |  |
| **SF-LV-shYIPF5_3-**<br>**mirE_rat** | This paper |  |
| **Cab45-eGFP**<br>**RUSH** | This paper |  |

### Cloning

Short hairpin RNAs (shRNAs) targeting rat *YIPF5* were designed based on the NM_001014150.3 reference sequence using the SPLASH RNA algorithm. The following sequences were selected and synthesized as 97 bp ultramers (IDT, USA): shRNA1:TCTATCAAAAACATCCTATCAT; shRNA2:TTAACAAACAAAACATTCCTAA; shRNA3: TTGTAAAGAAAATACCACTGCA Short hairpin RNAs targeting mouse *YIPF5* were designed based on the NM_023311.3 reference sequence using the same algorithm. The following sequences were synthesized as 97 bp ultramers (IDT, USA): shRNA1:TAAAGTAAGAAAATCATGTGTG; shRNA2:TTAACAAACAAAACATTCCTAA; shRNA3: TTGTAAAGAAAATACCACTGCA

For each shRNA, DNA fragments were amplified by PCR using GoTaq® G2 DNA Polymerase (Promega, USA) and specific primers (FWD_Gibson_5mirE-XhoI and REV_Gibson_3mirE-EcoRI). The resulting PCR products were cloned into the SF-LV-shRNA-mirE lentiviral vector, which drives shRNA expression under the control of the spleen focus-forming virus (SFFV) promoter, using NEBuilder® HiFi DNA Assembly Master Mix (New England Biolabs, USA). For each shRNA, three independent clones were sequenced using the FWD_EGFP-Ct primer (Eurofins Genomics) to verify correct insertion. YIPF5 disease-associated mutations (A181V, I98S, ΔL106, W218R) were introduced by site-directed mutagenesis using specific primer pairs and LeGO-iG2-Puro_hYIPF5 WT as templates. Reactions were performed with or without Pfu polymerase (Agilent) followed by PCR. PCR products were treated with DpnI for 3 h at 37 °C to digest template DNA, and mutagenized plasmids were validated by gel electrophoresis and sequencing. FLAG-hYIPF5 and V5-hYIPF5 inserts were ligated into BamHI/NotI-digested pcDNA3.1-hygro(+) using T4 DNA Ligase (NEB) at 1:3 and 1:5 vector:insert molar ratios, transformed into XL1-Blue cells by KCM protocol, and screened by restriction digest and sequencing. For shRNA expression, the lentiviral SGEP vector (Addgene #111170; gift from Johannes Zuber) was used. Three shRNAs targeting YIPF5 were designed: shRNA 1 (5′-TGC TGT TGA CAG TGA GCG ATA GGA ATG TTT TGT TTA TTA ATA GTG AAG CCA CAG ATG TAT TAA TAA ACA AAA CAT TCC TAG TGC CTA CTG CCT CGG A-3′), shRNA 2 (5′-TGC TGT TGA CAG TGA GCG CAG GAA TGT TTT GTT TAT TAA ATA GTG AAG CCA CAG ATG TAT TTA ATA AAC AAA ACA TTC CTA TGC CTA CTG CCT CGG A-3′), and shRNA 3 (′-TGC TGT TGA CAG TGA GCG AGA ACA CTT ACT TAC ATA TTA ATA GTG AAG CCA CAG ATG TAT TAA TAT GTA AGT AAG TGT TCG TGC CTA CTG CCT CGG A-3′). Oligos were amplified by PCR using Pwo polymerase (Roche) and ligated into EcoRI-HF/XhoI-digested SGEP by T4 DNA Ligase (NEB). The GCamPer coding sequence was amplified from Addgene plasmid #65227 (Henderson et al., 2015) using primers containing attB sites (forward: 5′- GGGGACAAGTTTGTACAAAAAAGCAGGCTTCCCGACTCTAGAGGATCC; reverse: 5′- GGGGACCACTTTGTACAAGAAAGCTGGGTCCTACGATAAGCTTGATATCGAATTC), gel-purified, and recombined with pDONR221 by BP Clonase™ II (Thermo Fisher Scientific). The pDONR221-GCamPer entry clone was verified and subsequently recombined with the pInd20 destination vector using LR Clonase™ II. For generation of the FUGW_Ii-Strep_L1CAM-APEX2-V5-SBP construct (referred to as L1CAM-V5 in this study), L1CAM was PCR-amplified from GW1-PAGFP-mL1CAM (gift from Casper Hoogenraad) using primers (forward: 5′- AAAACACGATGATAAgAATTATGGTCGTGATGCTGCGGTA; reverse: 5′- GCAGCaGAtCCAGCGGATCCTTCTAGGGCTACTGCAGGATTG). APEX2-V5-SBP was amplified from FUGW-RUSH-Ii-Strep_Nrxn1b(S4-)-APEX2-V5-SBP using primers (forward: 5′- GGATCCGCTGGATCTGCTGCAGGTTCTGGCGCTGGCTCCGCTGCTGGTTCT; reverse: 5′-GAATTGTTAACGGATCGAATTCTTATGGTTCACGTTGACCTTGTGG). FUGW_Ii-Strep was digested with EcoRI and BamHI, and L1CAM and APEX2-V5- SBP fragments were inserted by Gibson assembly. For generation of RUSH-Cab45- GFP, Cab45-HA ORF was cloned into the Str-KDEL_flGalT-SBP-EGFP plasmid (Addgene #65272) using the AscI site for the 3′ end and the EcoRI site for the 5′ end. CNPY3-mCherry expression plasmid was constructed in pCDH-MSCV-MCS-EF1α-41 Puro (Cat# CD710B-1, System Biosciences) by Gibson assembly as described in (Garloff et al., 2023).

### Mammalian tissue culture and transfection

All cell lines were maintained at 37 °C in a humidified incubator with 5% CO₂ under sterile conditions using a laminar flow hood. Subculturing was performed every 2–4 days upon reaching a maximum confluency of 80%, using trypsin (Sigma-Aldrich, cat. no. 59427C-100ML). HEK293T-LentiX (Takara Bio, cat. no. 632180), U2OS, NIH3T3 and HeLa cells were cultured in high-glucose DMEM supplemented with GlutaMax (Thermo Fisher Scientific, cat. no. 61965059), 10% fetal bovine serum (FBS, Thermo Fisher Scientific, cat. no. 10270106), and 1% penicillin-streptomycin (Thermo Fisher Scientific, cat. no. 15140122). MCF10A cells (ATCC CRL-10317) were maintained in DMEM/F-12 supplemented with 5% horse serum (Thermo Fisher Scientific, cat. no. 16050122), 1% penicillin-streptomycin, 10 μg/ml human insulin (Sigma-Aldrich, cat. no. I9278), 0.5 μg/ml hydrocortisone (Sigma-Aldrich, cat. no. H0888-5G), 0.1 μg/ml cholera toxin (Sigma-Aldrich, cat. no. C8052-2MG), and 20 ng/ml human epidermal growth factor (EGF, Biomol, cat. no. 50349.1000).

### Lentiviral production and transduction

For lentivirus production, HEK293T-LentiX cells were co-transfected with 10 µg psPAX2 (Addgene, #12260), 2.5 µg pMD2.G (Addgene, #12259), and 10 µg lentiviral plasmid using PEI-Max (Polysciences, cat. no. 24765-1). After 12 h, the medium was replaced, and lentiviral supernatants were collected at 36, 48, and 60 h post-transfection. Pooled supernatants were sterile-filtered and used to infect target cells for 24 h in culture medium supplemented with 8 µg/ml protamine sulfate (Sigma-Aldrich, cat. no. P4505-1G). Stable cell lines were generated through antibiotic selection. MCF10A YIPF5 KO cells (Cramer et al., 2023) were used to generate cells re-expressing YIPF5 wild type or one of the mutants.

For evaluating knockdown efficiency, primary cortical rat neurons or other target cells were transduced with lentiviruses to stably express the specific shRNA (100 µL of unconcentrated virus per well in a 12-well plate for neurons, and 1 mL per 10-cm dish for other cell types). For non-neuronal cells the efficiency was evaluated via WB.

### Transfection

For transient transfection, cells were seeded 24 h prior to transfection and incubated with a DNA–Lipofectamine™ 2000 (Thermo Fisher Scientific, cat. no. 11668019) complex, prepared according to the manufacturer’s protocol, using 2.4 µg DNA and 4.8 µl Lipofectamine per well.

### Primary hippocampal neuron culture

Hippocampi were dissected from embryonic day 18 (E18) rat brains, dissociated in trypsin for 15 min, and plated on poly-l-lysine (37.5 μg/ml) and laminin (1.25 μg/ml)- coated coverslips at a density of 50,000 or 100,000 cells per well in 12-well plates. The plating day was designated as day-in-vitro 0 (DIV0). Neurons were maintained in Neurobasal (NB) medium supplemented with 1% B27 (GIBCO), 0.5 mM glutamine (GIBCO), 15.6 μM glutamate (Sigma-Aldrich), and 1% penicillin-streptomycin (GIBCO) under standard incubation conditions (37 °C, 5% CO₂).

### Neuronal transfection

Primary hippocampal neurons were transfected using Lipofectamine 2000 (Invitrogen) according to the manufacturer’s instructions. Briefly, 0.05–2 μg of plasmid DNA was mixed with 1.2 μl of Lipofectamine 2000 in 200 μl Opti-MEM, incubated for 20 min at room temperature, and subsequently added to neurons in NB medium. After 1 h of incubation at 37 °C and 5% CO₂, neurons were washed with NB medium and returned to their original culture medium until fixation or imaging at the specified DIV time points.

### qPCR

RNA was extracted from cells cultivated in 6-well plates using the RNeasy Mini Kit from QIAGEN. Subsequently, cDNA was generated using the iScript™ cDNA Synthesis Kit from BioRad. The qPCR analysis in this study was conducted using the ViiA™ 7 Real-Time PCR System with a 384-Well Block from thermofisher.com. The primer mix was prepared by obtaining 100uM stock solutions of Forward (Fw) and Reverse (Rv) primers from the IDT company. The primers were then diluted to a working mix of 5uM Fw + 5uM Rv, which involved the combination of 10uL of 100uM Fw, 10uL of 100uM Rv, and 180uL of Molecular Biology Grade Water (MQ H2O). For each qPCR reaction (x1), the primer mix (F+R) was added at 0.5uL each, achieving a final concentration of 0.5uM for each primer. The reaction mix comprised of 1uL of cDNA, 1uL of MQ H2O, and 2.5uL of SYBR Green (ThermoScientific, 4309155). The total volume for each reaction was 5uL. The primer sequences for the target genes, GAPDH, b-actin, BIP and YIPF5, were as follows:

-Gapdh Forward primer: CAACTCCCTCAAGATTGTCAGCAA;

-Gapdh Reverse primer: GGCATGGACTGTGGTCATGA;

-b-actin Forward primer: AGGGAAATCGTGCGTGACAT;

-b-actin Reverse primer: ACGCAGCTCAGTAACAGTCC.

-YIPF5 Forward primer CCCCTTTGCTAGAAGAGTTGGG;

-YIPF5 Reverse primer GCCAAACTGGATTTTGCCCG.

### -BIP Forward primer GAAAGAAGGTTACCCATGCAGT

### -BIP Reverse primer CAGGCCATAAGCAATAGCAGC

The qPCR machine settings and parameters adhered to the specifications outlined by the ViiA™ 7 Real-Time PCR System manual.

### Co-Immunoprecipitation

Equal numbers of cells were cultured in 6-wells. The subsequent day, transfections were performed using the specified plasmids, following the manufacturer’s instructions for Lipofectamine 2000 (Thermo Fisher Scientific). After transfection, cells were lysed in CHAPSO lysis buffer (150 mM citrate buffer, pH 6.4, 2% CHAPSO) by scraping directly in the wells. Lysates were incubated on ice for 30 min, followed by centrifugation at maximum speed (≥13,000 × g) at 4 °C for 15 min. The supernatant was transferred to a fresh tube, and protein concentration was determined using the Pierce™ BCA Protein Assay Kit (Thermo Fisher Scientific, Cat. No. 23225).

For immunoprecipitation, GFP-TRAP agarose beads (ChromoTek) or anti-FLAG M2 agarose beads (Sigma-Aldrich) were equilibrated and washed extensively with 150 mM citrate buffer (pH 6.4). Equal amounts of protein lysates were incubated with pre-washed beads under continuous end-over-end rotation (14 rpm) at 4 °C overnight.

The following morning, beads were pelleted by centrifugation at 2,500 × g for 5 min at 4 °C and washed multiple times with 150 mM citrate buffer. Proteins were eluted by direct resuspension in 2× Laemmli buffer, followed by vigorous vortexing and boiling at 95 °C for 10 min with gentle shaking. After centrifugation at 2,500 × g for 5 min, the supernatant was collected and loaded onto an SDS-PAGE gel for protein analysis.

GFP-CAB45 was immunoprecipitated from the culture medium using GFP-Trap beads. HeLa cells stably expressing shScramble or shRNA3-YIPF5 were transfected with RUSH-GFP-CAB45 in 6 cm dishes. 4 hours after transfection, the medium was replaced, and cells were incubated overnight at 37 °C with 5% CO₂. 24 hours post-transfection, culture medium was replaced with 1 mL of fresh medium supplemented with biotin (final concentration: 40 µM) to synchronize secretion. Conditioned media were collected at the indicated time points post-biotin addition (0, 15, 30,60,120 and 180 minutes). Media were collected, supplemented with protease inhibitors and centrifuged to remove debris. A total of 900 µl of conditioned medium was incubated with 25 µL of GFP-Trap beads pre-equilibrated in 150 mM citrate buffer, overnight at 4 °C under rotation. Beads were washed twice with 150 mM citrate buffer + 0.5% CHAPSO and once with citrate buffer alone, then eluted with 2× Laemmli buffer and analyzed by SDS-PAGE and Western blot using an anti-GFP antibody.

### Endo H assay

Endo H digestion was performed in HeLa cells stably expressing either shScramble or shRNA3 targeting YIPF5. Cells were transiently transfected using Lipofectamine 2000 according to the manufacturer’s instructions. Transfections were carried out in the morning, followed by medium replacement 4 hours post-transfection. Cells were then incubated overnight at 37 °C with 5% CO₂ before the experiment. On the following day, cells were treated with 1ml/6cm dish prewarmed growth medium with 40 µg/ml Biotin (stock solution prepared at 40 mg/mL in DMSO) to synchronize cargo release from the ER. Media were collected at the indicated time points and cells were lysed in STEN buffer supplemented with protease inhibitors and total protein concentration was quantified. Equal protein amounts were subjected to Endo H digestion using the NEB Endo H kit. Briefly, 20 µL of each lysate sample was denatured at 95 °C for 10 minutes in glycoprotein denaturing buffer. Following denaturation, samples were incubated with Endo H enzyme at 37 °C for 2 hours in the provided reaction buffer. After digestion, Laemmli sample buffer was added to each reaction, and samples were boiled at 95 °C for 10 minutes.

### BCA-Assay

To determine the protein concentration of cell lysates, we used the Pierce bicinchoninic acid (BCA) Protein Assay Kit (Thermo Scientific™, cat.no. 23227) according to the manufacture instructions. The samplés absorbance was measured using a Mithras LB 940 multimode microplate reader. To determine the concentration of the samples, we referred to a standard curve created based on the absorbance of the enclosed bovine serum albumin.

### Stain-Free Gel Imaging

To estimate the total protein concentration within the polyacrylamide gels, a stain-free imaging technique was employed. The gels underwent a 5-minute incubation in 10% trichloroacetic acid (TCA), followed by a series of three washes with deionized water (dH2O). Visualization of the proteins was achieved using the UV illumination function on a Bio-Rad Gel Doc XR+ system.

### SDS-PAGE and Immunoblotting

Cells were cultured to confluence in 6-well plates or 10 cm dishes, rinsed twice with ice-cold PBS, and lysed in NP-40 lysis buffer (50 mM Tris–HCl, pH 7.6, 150 mM NaCl, 2 mM EDTA, 1% NP-40) supplemented with protease inhibitors. Following scraping and a 30-minute incubation on ice, lysates were centrifuged at 500 × g for 15 min at 4 °C, and the supernatant was collected for protein quantification using the Pierce™ BCA Protein Assay Kit (Thermo Fisher Scientific, Cat. No. 23225). Protein samples were diluted with 6× sample buffer (12% SDS, 0.0004% Bromophenol Blue, 47% glycerol, 12% Tris-HCl, pH 6.8, 9.3% DTT) and separated by SDS-PAGE. Proteins were transferred onto Immobilon-P PVDF membranes (Carl Roth, T831.1), with a pre-stained protein ladder (PageRuler™ Plus, Thermo Fisher, Cat. No. 26619) used as a molecular weight marker. Membranes were blocked with I-Block™ solution (1 g I- Block™ powder, 0.1% Tween-20 in 500 mL 1× PBS) for 1 h at room temperature, followed by incubation with primary antibodies diluted in blocking solution for 1 h at room temperature or overnight at 4 °C. After three washes (5 min each) with 1× TBS-Tween, membranes were incubated with horseradish peroxidase (HRP)-conjugated secondary antibodies for 30 min at room temperature. Following additional washes, protein signals were detected via chemiluminescence. For total protein visualization, stain-free gel imaging was performed prior to transfer by incubating gels in 10% (v/v) trichloroacetic acid for 5 min, followed by thorough washing with distilled deionized water (ddH2O).

### Proteomics

#### Sample Preparation for proteomics analysis

For whole cells analysis, cells were lysed in a buffer containing 5% SDS, 100 mM HEPES, and 50 mM DTT to ensure efficient protein extraction. The lysate was subjected to sonication using a Bioruptor Plus (Diagenode, Belgium) with 10 cycles of 30 seconds on and 60 seconds off, maintaining a temperature of 20°C to prevent protein degradation. Subsequently, 50 μg per sample samples were thermally denatured at 95°C for 5 minutes to fully solubilize proteins. Thiol groups were reduced and alkylated by incubating with 15 mM iodoacetamide in the dark at room temperature for 30 minutes. Proteins were precipitated overnight at -20°C after adding a 5x volume of ice-cold acetone. The following day, samples were centrifuged at 16,000 rcf and 4°C for 30 minutes, and the supernatant was carefully removed. Pellets were washed twice with 300 µL of ice-cold 80% (v/v) acetone in water, followed by centrifugation at 16,000 rcf and 4°C for 10 minutes. Pellets were air-dried before resuspension in 25 µL of digestion buffer (3 M urea, 100 mM HEPES, pH 8.0). Samples were resuspended using sonication (as described above), and LysC (Wako) was added at a 1:100 (w/w) enzyme-to-protein ratio. Digestion proceeded at 37°C for 4 hours with shaking (1000 rpm for 1 hour, then 650 rpm). Samples were then diluted 1:1 with Milli-Q water, and trypsin (Promega) was added at the same enzyme-to-protein ratio. Digestion continued overnight at 37°C with shaking (650 rpm). The following day, digests were acidified by adding trifluoroacetic acid (TFA) to a final concentration of 10% (v/v) and desalted using Waters Oasis® HLB µElution Plates (30 µm pore size, Waters Corporation) under a slow vacuum. Columns were conditioned with 3x 100 µL of solvent B (80% (v/v) acetonitrile, 0.05% (v/v) formic acid) and equilibrated with 3x 100 µL of solvent A (0.05% (v/v) formic acid in Milli-Q water). Samples were loaded, washed three times with 100 µL of solvent A, and eluted with 50 µL of solvent B into 0.2 mL PCR tubes. Eluates were dried using a speed vacuum centrifuge and reconstituted at a concentration of 1 µg/µL in reconstitution buffer (5% (v/v) acetonitrile, 0.1% (v/v) formic acid in Milli-Q water). Protein amounts were estimated based on cell number input and confirmed by SDS-PAGE analysis of 4% of each sample against an in-house cell lysate of known quantity. Reconstituted peptides were analyzed by data-independent acquisition (DIA).

### Secretome

For secretome analysis, first Streptavidin Sepharose High Performance beads (Cytiva) were acetylated to minimize non-specific binding. Beads were equilibrated in PBS and lysine-acetylated using 20 mM sulpho-NHS-acetate for 1 hour at room temperature. The reaction was quenched by adding 1 M Tris pH 7.5, and beads were extensively washed with PBS to remove unreacted reagents. Glycoprotein enrichment from the cellular secretome was achieved using a metabolic labeling approach involving click sugars (Serdaroglu et al., 2017; Tüshaus et al., 2020). Cells from five 10 cm plates per experimental condition were cultured in the presence of 62.5 µM Click-IT™ ManNAz Metabolic Glycoprotein Labeling Reagent (Invitrogen) for 48 hours to label secreted glycoproteins. Conditioned media were supplemented with EDTA-free protease inhibitors to prevent proteolysis and filtered through a 0.45 µm filter. Media volumes were normalized to account for equal cell numbers and concentrated using Vivaspin 20 concentrators with a 30 kDa molecular weight cutoff by centrifugation at 4000×g for 80 minutes at 4°C, reducing the volume to 0.5 mL. Concentrated samples were washed twice with 15 mL of PBS at 4°C to remove unbound molecules and spent media. Final volumes were adjusted to 1 mL, and 100 µM DBCO-Sulfo-Biotin (Jena Bioscience, Germany) was added for overnight incubation at 4°C to enable click chemistry-mediated biotinylation.

For glycoprotein isolation, Concanavalin A (Con A) agarose beads (Sigma-Aldrich) were prepared by washing twice with 1 mL of binding buffer (20 mM Tris-HCl pH 7.5, 500 mM NaCl, 5 mM MgCl2, MnCl2, and CaCl2). Samples were mixed with 300 µL of prepared Con A beads and 1 mL of binding buffer in low-binding tubes and incubated at 4°C with rotation for 2 hours to permit glycoprotein binding. Post-incubation, beads were washed three times with binding buffer to remove non-specific binders. Glycoproteins were eluted from the beads with two rounds of 500 µL elution buffer (20 mM Tris-HCl pH 7.5, 500 mM Methyl-α-D-mannopyranoside, 10 mM EDTA) at 4°C for 30 minutes each. Eluates were combined and passed through Pierce™ Spin Columns (Thermo Fisher Scientific) to remove agarose beads. For each sample 300 µl of acetylated streptavidin beads were equilibrated with PBS before adding ConA enriched samples and incubated overnight at 4 °C rotating at 15 rpm. Afterwards, samples were centrifuged for 5 min at 4 °C and 2000×g, and beads were resuspended in PBS and transferred to a Spin Column. Beads were washed once with 1 ml PBS, three times with 1 ml wash buffer 1 (30 mM AmBic, 3M Urea) and finally 5 times with 600 µl wash buffer 2 (50 mM AmBic pH 8.0). These serial washes were designed to rigorously remove non-specific proteins before downstream proteomic analysis.

### Surfaceome

For surfaceome analysis, cells were cultured on 10 cm dishes for 48 hours (up to 20 million cells), with all subsequent procedures performed on ice to maintain protein integrity. Cells were washed three times with 5 mL of ice-cold PBS supplemented with calcium and magnesium, followed by incubation with 3 mL of 0.5 mg/mL EZ-Link™ Sulfo-NHS-LC-Biotin (Thermo Fisher Scientific) for 30 minutes to biotinylate surface-exposed proteins. Unreacted biotinylation reagent was quenched by washing cells four times with 5 mL of 20 mM glycine in PBS for 15 minutes each, followed by a final wash with PBS containing calcium and magnesium. Cells were then lysed with surface specific lysis buffer (50 mM Tris, 150 mM NaCl, 2mM EDTA, 0,2% NP40). For each sample 80 µl of acetylated beads were equilibrated in surface lysis buffer before lysates were added overnight at 4°C rotating at 15 rpm. Beads were washed with 1 ml wash buffer 3 (50mM Tris-HCl pH 7.6, 325mM NaCl, 2mM EDTA, 0.2% NP-40), 4 ml wash buffer 4 (50mM Tris-HCl pH 7.6, 150mM NaCl, 2mM EDTA, 1% NP-40, 1% SDS) and 5 ml wash buffer 5 (50mM Tris-HCl pH 7.6, 150mM NaCl, 2mM EDTA). The beads were finally washed 5 times with 600 µl wash buffer 2 (50 mM AmBic pH 8.0). to remove non-specifically bound proteins.

For both procedures, beads were transferred to a fresh tube using three times 300 μl wash buffer, centrifuged at 2000×g for 5 min at 4 °C and resuspended in 200 μl wash buffer 2 and 1 µg LysC added. After an incubation overnight at 37°C, peptides were eluted with two times 150µl wash buffer 2. Elutions were further digested with 0.5 µg trypsin for 3 hours at 37°C. The day after, digests were acidified by the addition of TFA to a final concentration of 1% (v/v), then desalted with Waters Oasis® HLB µElution Plate 30 µm (Waters Corporation, MA, USA) under a soft vacuum following the manufacturer instruction. Briefly, the columns were conditioned with 3x100 µL solvent B (80% (v/v) acetonitrile; 0.05% (v/v) formic acid) and equilibrated with 3x100 µL solvent A (0.05% (v/v) formic acid in Milli-Q water). The samples were loaded, washed 3 times with 100 µL solvent A, and then eluted into 0.2 mL PCR tubes with solvent B. Samples were dried with a speed vacuum centrifuge and stored at −20 °C until LC-MS analysis.

### Data independent acquisition (DIA) and data analysis

For whole cell, samples were reconstituted in in MS Buffer (5% acetonitrile, 95% Milli-Q water, with 0.1% formic acid) and spiked with iRT peptides (Biognosys, Switzerland). Peptides were separated in trap/elute mode using the nanoAcquity MClass Ultra-High Performance Liquid Chromatography system (Waters, Waters Corporation, Milford, MA,USA) equipped with a trapping (nanoAcquity Symmetry C18, 5 μm, 180 μm × 20 mm) and an analytical column (nanoAcquity BEH C18, 1.7 μm, 75 μm × 250 mm). Solvent A was water and 0.1% formic acid, and solvent B was acetonitrile and 0.1% formic acid. 1 µl of the sample (∼1 μg on column) were loaded with a constant flow of solvent A at 5 μl/min onto the trapping column. Trapping time was 6 min. Peptides were eluted via the analytical column with a constant flow of 0.3 μl/min. During the elution, the percentage of solvent B increased in a nonlinear fashion from 0–40% in 120 min. Total run time was 145 min. including equilibration and conditioning. The LC was coupled to an Orbitrap Exploris 480 (Thermo Fisher Scientific, Bremen, Germany) using the Proxeon nanospray source. The peptides were introduced into the mass spectrometer via a Pico-Tip Emitter 360-μm outer diameter × 20-μm inner diameter, 10-μm tip (New Objective) heated at 300 °C, and a spray voltage of 2.2 kV was applied. The capillary temperature was set at 300°C. The radio frequency ion funnel was set to 30%. For DIA data acquisition, full scan mass spectrometry (MS) spectra with mass range 350–1650 m/z were acquired in profile mode in the Orbitrap with resolution of 120,000 FWHM. The default charge state was set to 3+. The filling time was set at maximum of 60 ms with limitation of 3 × 10^6^ ions. DIA scans were acquired with 40 mass window segments of differing widths across the MS1 mass range. Higher collisional dissociation fragmentation (stepped normalized collision energy; 25, 27.5, and 30%) was applied and MS/MS spectra were acquired with a resolution of 30,000 FWHM with a fixed first mass of 200 m/z after accumulation of 3 × 10^6^ ions or after filling time of 35 ms (whichever occurred first). Data were acquired in profile mode. For data acquisition and processing of the raw data Xcalibur 4.3 (Thermo) and Tune version 2.0 were used.

For surfeacome and secretome analysis, samples were reconstituted in in MS Buffer (5% acetonitrile, 95% Milli-Q water, with 0.1% formic acid) and spiked with iRT peptides (Biognosys, Switzerland). Peptides were separated in trap/elute mode using the nanoAcquity MClass Ultra-High Performance Liquid Chromatography system (Waters, Waters Corporation, Milford, MA, USA) equipped with a trapping (nanoAcquity Symmetry C18, 5 μm, 180 μm × 20 mm) and an analytical column (nanoAcquity BEH C18, 1.7 μm, 75 μm × 250 mm). Solvent A was water and 0.1% formic acid, and solvent B was acetonitrile and 0.1% formic acid. 1 µl of the sample (∼1 μg on column) were loaded with a constant flow of solvent A at 5 μl/min onto the trapping column. Trapping time was 6 min. Peptides were eluted via the analytical column with a constant flow of 0.3 μl/min. During the elution, the percentage of solvent B increased in a nonlinear fashion from 0–40% in 90 min. Total run time was 120 min. including equilibration and conditioning. The LC was coupled to an Orbitrap Exploris 480 (Thermo Fisher Scientific, Bremen, Germany) using the Proxeon nanospray source. The peptides were introduced into the mass spectrometer via a Pico-Tip Emitter 360-μm outer diameter × 20-μm inner diameter, 10-μm tip (New Objective) heated at 300 °C, and a spray voltage of 2.2 kV was applied. The capillary temperature was set at 300°C. The radio frequency ion funnel was set to 30%. For DIA data acquisition, full scan mass spectrometry (MS) spectra with mass range 350–1650 m/z were acquired in profile mode in the Orbitrap with resolution of 120,000 FWHM. The default charge state was set to 3+. The filling time was set at maximum of 60 ms with limitation of 3 × 10^6^ ions. DIA scans were acquired with 34 mass window segments of differing widths across the MS1 mass range. Higher collisional dissociation fragmentation (stepped normalized collision energy; 25, 27.5, and 30%) was applied and MS/MS spectra were acquired with a resolution of 30,000 FWHM with a fixed first mass of 200 m/z after accumulation of 3 × 10^6^ ions or after filling time of 35 ms (whichever occurred first). Data were acquired in profile mode. For data acquisition and processing of the raw data Xcalibur 4.3 (Thermo) and Tune version 2.0 were used. For IP analysis, peptides were separated using the Evosep One system (Evosep, Odense, Denmark) equipped either with a 15 cm x 150 μm i.d. packed with a 3 μm Reprosil-Pur C18 bead column (Evosep Endurance, EV-1109, PepSep, Marslev, Denmark) or with a a 15 cm x 150 μm i.d. packed with a 1.5 μm Reprosil-Pur C18 bead column (Evosep Performance, EV-1137, PepSep, Marslev, Denmark) heated at 45°C with a butterfly sleeve oven (Phoenix S&T, Philadelphia, USA). The samples were run with a pre-programmed proprietary Evosep gradient of 44 min (30 samples per day) using water and 0.1% formic acid and solvent B acetonitrile and 0.1% formic acid as solvents. The LC was coupled to an Orbitrap Exploris 480 (Thermo Fisher Scientific, Bremen, Germany) using PepSep Sprayers and a Proxeon nanospray source. The peptides were introduced into the mass spectrometer via a PepSep Emitter 360-μm outer diameter × 20-μm inner diameter, heated at 300°C, and a spray voltage of 2 kV was applied. The injection capillary temperature was set at 300°C. The radio frequency ion funnel was set to 30%. For DIA data acquisition with endurance column, full scan mass spectrometry (MS) spectra with a mass range of 350–1650 m/z were acquired in profile mode in the Orbitrap with a resolution of 120,000 FWHM. The default charge state was set to 2+, and the filling time was set at a maximum of 60 ms with a limitation of 3 × 10^6^ ions. DIA scans were acquired with 40 mass window segments of differing widths across the MS1 mass range. Higher collisional dissociation fragmentation (normalized collision energy 29%) was applied, and MS/MS spectra were acquired with a resolution of 30,000 FWHM with a fixed first mass of 200 m/z after accumulation of 1 × 10^6^ ions or after filling time of 45 ms (whichever occurred first). Data were acquired in profile mode. For data acquisition and processing of the raw data, Xcalibur 4.5 (Thermo Fisher) and Tune version 4.0 were used.

### Data Analysis for DIA Samples

For whole cell analysis, DIA raw data were analyzed using the directDIA pipeline in Spectronaut (v. 14.9, Biognosys AG). The data were searched against a species-specific SwissProt database tailored for *Homo sapiens*, comprising 20,816 protein entries, as well as a contaminant database with 247 entries, including the following modifications: Oxidation (M), Acetyl (Protein N-term). A maximum of 2 missed cleavages for trypsin and 5 variable modifications were allowed. Local normalization according to (Callister et al., 2006) was applied. The identifications were filtered to satisfy FDR of 1 % on peptide and protein level. Relative quantification was performed in Spectronaut for each paired comparison using the replicate samples from each condition.

For secretome and surfaceome analysis, DIA raw data were analyzed using the directDIA pipeline in Spectronaut (Biognosys AG, v. 16 for surfaceome and secretome, v. 18 for IPs,). The data were searched against a species-specific SwissProt database tailored for *Homo sapiens*, comprising 20,816 protein entries, as well as a contaminant database with 247 entries, including the following modifications: Oxidation (M), Acetyl (Protein N-term). A maximum of 2 missed cleavages for trypsin and 5 variable modifications were allowed. Local normalization according to Callister et al (2006) was applied. The identifications were filtered to satisfy FDR of 1 % on peptide and protein level. Relative quantification was performed in Spectronaut for each paired comparison using the replicate samples from each condition. Precursor were filtered if identified in at leats 20% of the runs and a global imputation was applied. Data processing and statistical analyses were conducted using customized R scripts executed in RStudio. As a primary pre-processing step for both secretome and surfaceome data, a stringent filtering protocol was applied to remove potential intracellular contaminants. To achieve this, we first parsed protein localization information from (Binder et al., 2014). Any protein exhibiting a maximum localization score of 5 for either the “nucleus” or “mitochondrion” compartments was filtered out. An additional filtering step was performed to remove proteins associated with the Gene Ontology term “ribosome” (GO:0005840). The resulting clean datasets were then used for all downstream statistical analyses. Significant protein abundance changes were using a q-value threshold of <0.05 for all experiments except for the secretome and surfaceome analyses, where the q-value was set to <0.1. Protein in surfaceome and secretome data were marked as “explained by proteome” if the direction of the change was congruent with the one observed at the proteome level.

### Data availability

The proteomic datasets generated during and/or analyzed during the current study are available in the MassIve repository, secretome dataset ID MSV000093769, surfaceome dataset ID MSV000093771, total proteome dataset ID MSV000093778.

### Immunofluorescence

Non-neuronal cells were cultured on glass coverslips (Paul Marienfeld GmbH, Germany, Cat. No. 0111550) and subjected to experimental treatments as indicated. Fixation was performed at room temperature (RT) for 20 min using ice-cold 4% paraformaldehyde diluted in PBS, followed by three washes with 1× PBS (5 min each). Permeabilization was carried out with 0.2% Triton X-100 in PBS (RT, 5 min), and non-specific binding sites were blocked using a solution containing 1 ml fetal calf serum (FCS), 1 g bovine serum albumin (BSA), and 0.1 g fish gelatin in 100 ml PBS for 20 min at RT. Cells were incubated with primary antibodies diluted in blocking solution for 1 h at RT, followed by three washes (5 min each) with PBS. Subsequently, samples were incubated with Alexa Fluor 488-, 555-, or 647-conjugated secondary antibodies (1:500 dilution, Invitrogen) for 30 min at RT. Following incubation, cells were washed and stained with Hoechst 33342 (1:10,000 dilution in PBS, Sigma-Aldrich, Cat. No. B2261-25MG) for 10 min. After three additional washes with warm PBS, coverslips were mounted on microscope slides (Epredia, Germany, Cat. No. AA00000112E01MNZ10) using Mowiol 4–88. Imaging was performed using a Zeiss Imager Z.2 microscope with an Apotome.2 structured illumination system, an Axiocam 702 camera, and a Plan-Apochromat 63×/1.4 Oil DIC M27 objective (Carl Zeiss AG, Germany) or a Zeiss LSM 880 microscope with an AiryScan detector, utilizing a 488 nm argon laser (Melles-Griot) and a Plan-Apochromat 63×/1.4 NA DIC M27 oil objective.

Neurons were fixed at RT for 20 min with 4% paraformaldehyde, followed by permeabilization with 0.2% Triton X-100 in PBS supplemented with calcium and magnesium (PBS-CM) for 15 min. Blocking was performed at 37 °C for 30 min using 0.2% porcine gelatin in PBS-CM. Neurons were then incubated with primary antibodies, followed by secondary antibody incubation (30 min, RT). After antibody incubations, cells were washed three times (5 min each) with PBS-CM. DAPI (0.1 µg/ml in PBS) was applied for 5 min for nuclear staining, followed by additional PBS-CM washes. Coverslips were mounted using Fluoromount-G Mounting Medium (Thermo Fisher Scientific). Images were acquired using a Zeiss LSM700 confocal laser-scanning microscope, equipped with Zen imaging software (version 8.1.7.484), a Plan-Apochromat 63×/1.4 NA oil DIC objective, and an EC Plan-Neofluar 40×/1.3 NA oil DIC objective.

For some experiments an ImageXpress Micro Confocal microscope (Molecular Devices, USA) equipped with a 60x Lambda air immersion objective, temperature and humidity control (Molecular Devices, CA), Air/CO2 gas mixer from OKOLAB, Italy, and MetaXpress High-Content Image Acquisition and Analysis v.6.7.2.290 (64-bit) software was used. For this, cells were cultivated in a 96-Well Cell Imaging Plate with a cover glass bottom (Eppendorf, Cat. No. 0030 741.030). Fixation was performed at RT for 10 min using 4% formaldehyde (Thermo Fisher Scientific, Cat. No. 28906) diluted in PBS, followed by multiple washes with 1× PBS (5 min each). Cells were then permeabilized with 0.2% Triton X-100 in PBS at RT for 5 min, washed again with PBS, and blocked with 4% BSA (Sigma-Aldrich, Cat. No. 232-936-2) in PBS at RT for 30 min. Primary antibody incubation was carried out at RT for 1 h, with antibodies diluted in blocking buffer (FCS, BSA, fish gelatin in PBS). Samples were subsequently incubated with Alexa Fluor 488-, 555-, or 647-conjugated secondary antibodies (1:1000 dilution, Invitrogen) for 60 min at RT, followed by nuclear staining with Hoechst 33342 (1:10,000 dilution) for 10 min. After three washes with warm PBS, fluorescence measurements were acquired.

### Live imaging

For live imaging, cells were cultured in a 96-Well Cell Imaging Plate with a cover glass bottom (Eppendorf, Cat. No. 0030 741.030). The media were replaced with DMEM high glucose HEPES without phenol red, and 5 µg/ml Hoechst 33342 prior to imaging. For ER-Ca^2+^ analysis, cells were plated in complete MCF10A media supplemented with 50mg/ml doxycycline one day before the acquisition. Half an hour before imaging, the media was replaced with HBSS without phenol red and 5 µg/ml Hoechst 33342. Cells were imaged using the ImageXpress microscope.

### Airyscan live-Cell Imaging

For super resolution live-cell imaging, cells were plated in MatTek glass-bottom dishes (Cat. No. P35G-1.5-14-C NS) and imaged using a Zeiss LSM 880 microscope equipped with an AiryScan detector and a Zeiss Plan-Apochromat 63×/1.4 NA DIC M27 oil objective. Excitation was achieved using an argon laser (Melles-Griot) at 488 nm. 37 °C and 5% CO₂ were maintained using the integrated temperature and CO₂ control system.

### ER-shape analysis

U2OS cells were seeded in 10 cm dishes to reach 70–90% confluency and transfected with the mRFP-KDEL plasmid (Snapp et al., 2006) using Lipofectamine™ 2000, following the manufacturer’s protocol. The medium was replaced after 6 hours, and RFP expression was confirmed by fluorescence microscopy the next day. Two days post-transfection, selection was initiated with complete DMEM supplemented with 100 μg/mL Hygromycin B. Medium was changed every 2 days. Selection continued until all control cells were dead. Surviving cells were subjected to limiting dilution (3 cells/100 μL) in 96-well plates using selective medium. Clones were monitored for 3 weeks, and positive monoclonal populations were expanded and validated for stable mRFP-KDEL expression.

U2OS-mRFP-KDEL-shScramble as control and U2OS-mRFP-KDEL-shRNA2-YIPF5 and U2OS-mRFP-KDEL-shRNA3-YIPF5 were generated by infecting U2OS with lentiviral SGEP-shRNAs as described above. Imaging for this assay was conducted in the 96-Well Cell Imaging Plates using the ImageXpress microscope as described above. We use the Custom Module Editor from MetaXpress software (Molecular Devices) to analyze the collected image data. Our image analysis algorithm comprises sequential processing steps: (i) detection of cell nuclei, (ii) segmentation of the cellular area, and (iii) implementation of the “Find Fibers” function. The initial phase utilizes Hoechst staining outcomes (acquired from the blue channel) for nuclear detection, laying the groundwork for subsequent cell-specific analysis. This step also aids in defining the cell territory for subsequent ER network analysis. The secondary phase generates an expanded mask around each nucleus to approximate the overall cellular domain, which captures the mRFP-tagged ER network within individual cells. The tertiary phase deploys the “Find Fibers” module that detects and segments fiber-like structures, in this case, the mRFP-KDEL labeled ER tubules, rendering a skeletonized binary representation for each cell analyzed. The ER’s complex morphology is characterized using distinct parameters extracted from the fiber segmentation mask. We focus on two metrics indicatives of the ER network’s intricacy: the total fiber count and the number of branch points. These chosen features provide a quantitative assessment, differentiating between normal and aberrant ER structures.

### IncuCyte cell proliferation assay

To assess cell proliferation in real time, cells were seeded at a density of 1.5 × 10⁴ cells/cm² in 96-well plates containing 100 µL of complete growth medium per well. Following seeding, live-cell imaging was initiated (t = 0 h) using the IncuCyte® SX5 Live-Cell Analysis System (Essen BioScience). Image acquisition was performed every 3 hours over a 72-hour period, capturing four non-overlapping fields per well using a 10× objective.For each experimental condition, three technical replicates (three wells per condition) were included. Cell confluence, representing the area occupied by cells, was quantified using the AI Confluence Analysis module within the IncuCyte software. Confluence values obtained from the four images per well were averaged to generate a single output per well.

### Incucyte scratch wound assay

A consistent and controlled wound model was established using the Incucyte® 96-Well Wound maker Tool to assess cell migration and wound healing. Monolayers of cells were plated at a density of 40,000 cells per well onto Incucyte® Imagelock 96-Well Plates (Sartorius, BA-04855) and allowed to settle and adhere during an over-night incubation period in a standard cell culture incubator. Subsequently, these confluent monolayers were uniformly wounded across all wells using the designated wound maker tool, producing wounds that were approximately 700–800 micrometers in width. Immediately after wound induction, the pre-used media was carefully aspirated from each well. The wounded monolayers were washed twice with fresh culture media, a step taken to prevent cell fragments from re-adhesion and potential interference with the wound healing observations. Fresh media was introduced to each well with a final volume of 100 μl. Before imaging, any air bubbles were removed from the wells to prevent inconsistencies in image analysis. After that, the plate was transferred to the Incucyte® Live-Cell Analysis System. A 5-minute equilibration period was allowed before initiating imaging to ensure optimal conditions for cell monitoring. The system was then programmed to scan each well hourly over 48 hours automatically. The live-cell analysis software was set up accordingly: “Scratch Wound” was selected as the scan type, and the “Wide Mode” option was chosen given the use of 10X objective lenses.

### In Utero Electroporation

In utero electroporation of embryonic mouse brains was performed following the protocol established by Mestres and Calegari (Mestres & Calegari, 2023) under the approved animal license TVV 16/2018 (DD24.1-5131/449/18; 25.05.2018-24.05.2023) and TVV 49/2022 at CRTD Zentrum für Regenerative Therapien TU Dresden. Timed-pregnant C57BL/6J mice at embryonic day 13.5 (Janvier Labs) were anesthetized using isoflurane.

Approximately 2 μl of a solution containing 2 mg/ml of plasmid mixed with 0.01% Fast Green dye was microinjected into the lateral ventricles of embryonic brains using a pulled glass capillary. The embryos were subsequently subjected to electroporation using an ECM830 Square Wave Electroporation System (BTX, MA), applying six pulses of 30 volts for 5 milliseconds with 950-millisecond intervals. Following electroporation, the uterine horns were repositioned, and the surgical incisions were sutured. The pregnant mice were allowed to recover in a warmed chamber before being returned to their home cages.

Two days post-electroporation, embryos were collected, and their brains were extracted and fixed in 4% paraformaldehyde overnight at 4°C. After fixation, 40 μm coronal sections were obtained using a Microm HM 650 V Vibratome (Thermo Fisher Scientific) and mounted onto Superfrost™ Plus Adhesion Microscope Slides (J1800AMNZ, Epredia, Germany). The slides were air-dried for 30 minutes at room temperature and subsequently washed in PBS before undergoing antigen retrieval in 10 mM sodium citrate buffer (pH 6.0) using a microwave heating protocol.

For immunohistochemistry, brain sections were incubated in blocking solution (5% normal goat serum, 1% BSA, and 0.1% Triton X-100 in PBS) for 2 hours at room temperature in a humidified chamber. Primary antibodies were diluted in blocking solution and incubated overnight at 4°C. The next day, sections were washed and incubated with secondary antibodies (1:1000 dilution) for 2 hours at room temperature. To reduce autofluorescence, sections were treated with 0.1% Sudan Black B in 70% ethanol for 20 minutes, followed by washes in PBS containing 0.02% Tween. Finally, the sections were mounted using Fluoromount™ Aqueous Mounting Medium (#F4680, Sigma-Aldrich) and sealed with #1 coverslips (H878 and 1870.2, Carl Roth).

Images were acquired using an Imager Z.2 with an Apotome.2 and an Axiocam 702, equipped with a Plan-Apochromat 63x/1.4 Oil DIC M27 objective (Carl Zeiss AG). Maximal intensity projections of serial Z-stacks were analyzed using Fiji software. Neuronal morphology in the cortical plate was assessed based on EGFP expression, and cells were classified as unbranched uni/bipolar or complex, based on the leading process structure.

### Statistics

All statistical tests were performed using R or GraphPad Prism. The specific statistical test and number of biological replicates are always indicated in the respective figure legend.

## Acknowledgements

We are grateful to the Core Facilities at the Fritz Lipmann Institute (FLI), Jena, Germany, for their invaluable support, including the Imaging Facility, the Core Facility Functional Genomics (Torsten Kroll), the Flow Cytometry (FACS) Facility. We especially thank Daniela Reichenbach, Jana Hamann, and Lea Hartmann for their excellent technical assistance, Johanna Mayer and Paul Cramer for their contribution to the generation of the YIPF5-KO and disease mutant cell lines. We are thankful to the SAND Innovative Training Network (ITN) for the stimulating scientific discussions and insightful input, with special thanks to Prof. Dr. Christian Behrends for his valuable feedback and guidance. CK received funding from the European Union’s Horizon 2020 research and innovation program under the Marie Skłodowska-Curie grant agreement No. 860035 (SAND) and from the DFG, grant #KA1751/8-1. JM and EAM were supported by the Wellcome Trust (225216/Z/22/Z) and by the Medical Research Council as part of United Kingdom Research and Innovation (MC_UP1201/10). For the purpose of open access, the MRC Laboratory of Molecular Biology has applied a CC BY public copyright license to any Author Accepted Manuscript version arising. FC was supported by CRTD Dresden, the School of Medicine of the TU Dresden, and the DFG, grant numbers: CA 893/9-1 and GRK2773/1-454245598.

## Author contributions

FB, MA, WD, MS conducted experiments and analyzed data; DDF analyzed data; TD, EC, AO conducted MS-related work and analysis; JM and EAM cloned plasmids and generated cell lines; IR and VG cloned plasmids; NK, GF cloned plasmids and provided rat hippocampal cultures; IM, FC and CV conducted and analyzed in-utero electroporation experiments. FB and CK conceived the study and wrote the manuscript with help of all authors. All authors read and approved the final version of the manuscript.

---

**Suppl. Figure 1.**
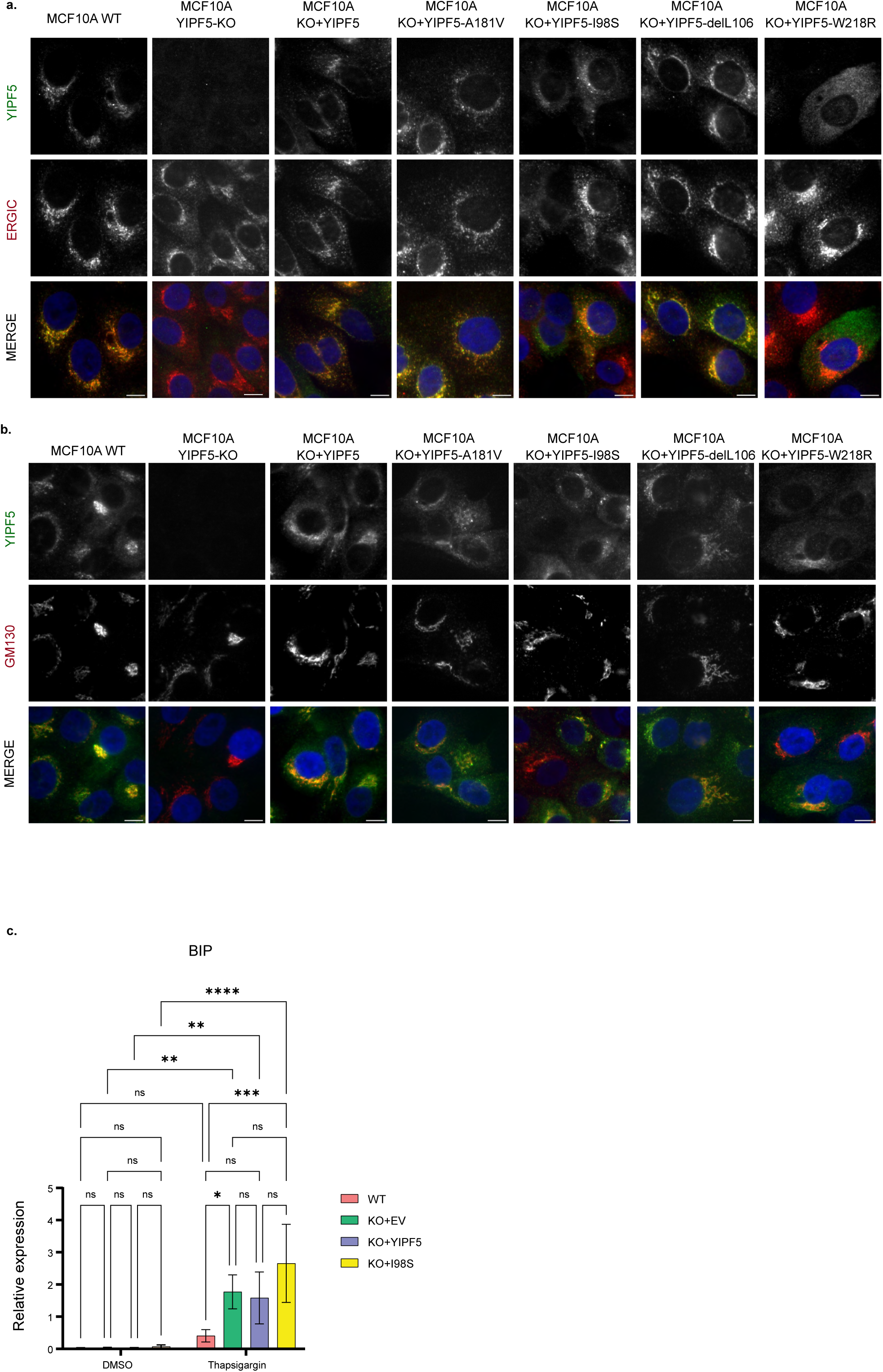
YIPF5 localizes to the ERGIC and Golgi apparatus and modulates ER stress response in MCF10A cells. MCF10A, MCF10A YIPF5-KO, and MCF10A YIPF5-KO cells re-expressing YIPF5 variants were fixed and (a) labeled with anti-YIPF5 (green) and anti-ERGIC-53 (red) antibodies or (b) labeled with anti-YIPF5 (green) and anti-GM130 (red) antibodies. Nuclei were stained with Hoechst dye (blue). Images were acquired using ImageXpress High-content confocal fluorescence microscopy. Scale bars: 10 µm. (c) Quantitative PCR analysis of BIP mRNA levels in MCF10A, MCF10A YIPF5-KO, and MCF10A YIPF5-KO cells re-expressing YIPF5 or the I98S variant. Cells were treated with DMSO or Thapsigargin (1µl/mL for 18h) to induce ER stress. Data are presented as mean ± SD. Statistical analysis was performed using two-way ANOVA with Šídák’s multiple comparisons test. (n = 3; * indicates P <0.05, *** indicates p < 0.001, **** indicates P < 0.0001).

**Suppl. Figure 2.**
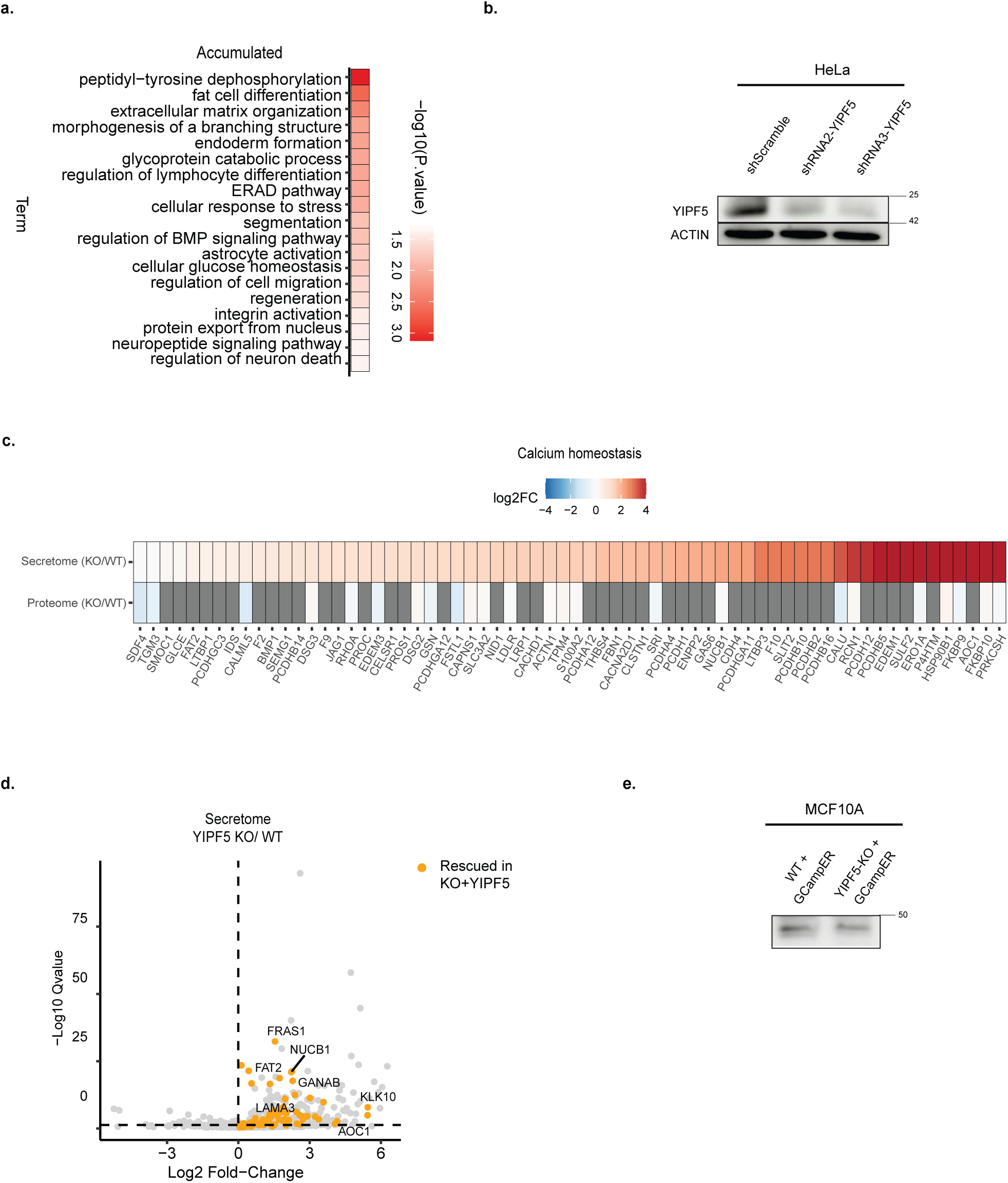
YIPF5 regulates calcium-related protein secretion and rescues dysregulated secretory pathways in MCF10A KO cells. (a) GO enrichment analysis of proteins with increased secretion in YIPF5 KO MCF10A cells compared to wild-type controls. Statistically significant categories are highlighted in red, demonstrating the broad impact of YIPF5 deficiency on secretory pathways and cellular processes. (b) Western blot analysis of HeLa cell lines stabling expressing shScramble, shRNA2-YIPF5 or shRNA3-YIPF5 using anti-YIPF5 and anti-Actin (loading control) (c) Heatmap displaying proteins associated with calcium ion transport (GO:0006816) and calcium ion binding (GO:0005509) that are differentially expressed in the secretome of YIPF5 KO MCF10A cells compared to WT, grey color indicates protein not detected at the proteome level. (d) Analysis of protein abundance changes in the secretome of YIPF5 KO cells re-expressing YIPF5 compared to YIPF5-KO cells. Proteins whose secretion is significantly restored by YIPF5 re-expression are highlighted in orange. The vertical axis represents -log10(q-value), and the horizontal axis represents log2(fold change). (e) Western blot analysis of MCF10A WT and YIPF5-KO cell lines stabling expressing GCampER using anti-GFP. Source data are available for this figure: SourceData FS2.

**Suppl. Figure 3.**
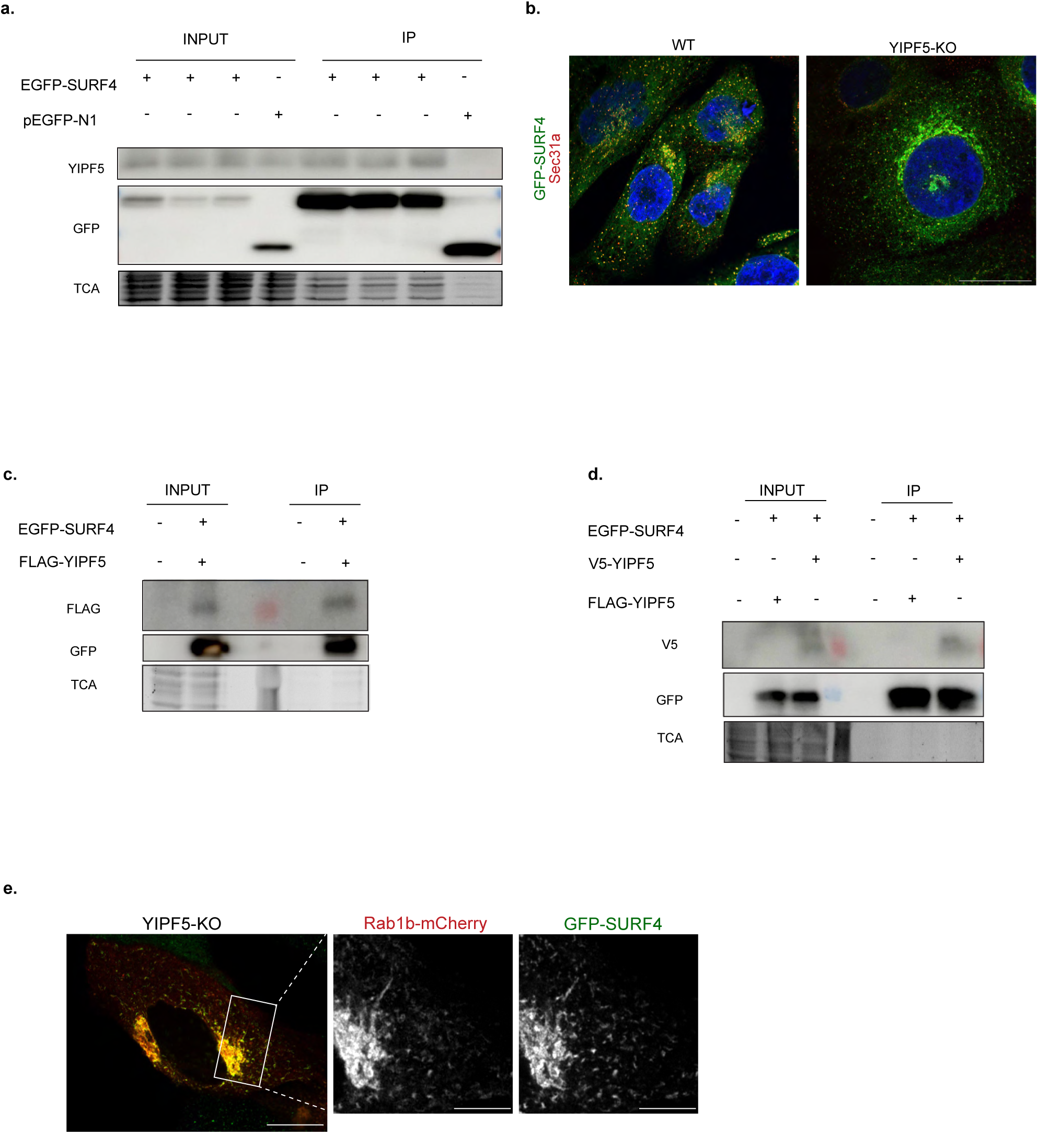
SURF4 interacts and co-localizes with YIPF5. (a) Cells were transiently transfected with EGFP-SURF4 and either FLAG-YIPF5, V5-YIPF5, or EGFP (negative control), lysed after 24 h, immunoprecipitated using GFP-TRAP beads and analyzed by immunoblotting with the indicated antibodies. Total lysates and the IP fractions are shown. TCA shows protein loading across conditions. (b) Airyscan super-resolution full images of Fig. 3d, illustrating SURF4 and SEC31A co-staining in YIPF5 knockout (KO) cells. Scale bar: 10 µm. (c) Independent Co-IP validation of the SURF4–YIPF5 interaction. HeLa cells were transfected with or without EGFP-SURF4 and FLAG-YIPF5, followed by GFP-TRAP pull-down and Western Blotting as described in (a). (d) Co-IP of EGFP-SURF4 with either V5-YIPF5 or FLAG-YIPF5, including an untransfected control. Binding was assessed via GFP-TRAP followed by immunoblotting of IP and control fractions as in (a) and (c). (e) Single frame from live-cell Airyscan super-resolution imaging of YIPF5-KO MCF10A cells stably expressing GFP-SURF4 (green) and transiently transfected with Rab1b-mCherry (red). Source data are available for this figure: SourceData FS3. The full movie is shown in video 3. Rab1b associates with SURF4-positive tubular structures. Scale bars: overview, 10 µm; insets, 2 µm.

**Suppl. Figure 4.**
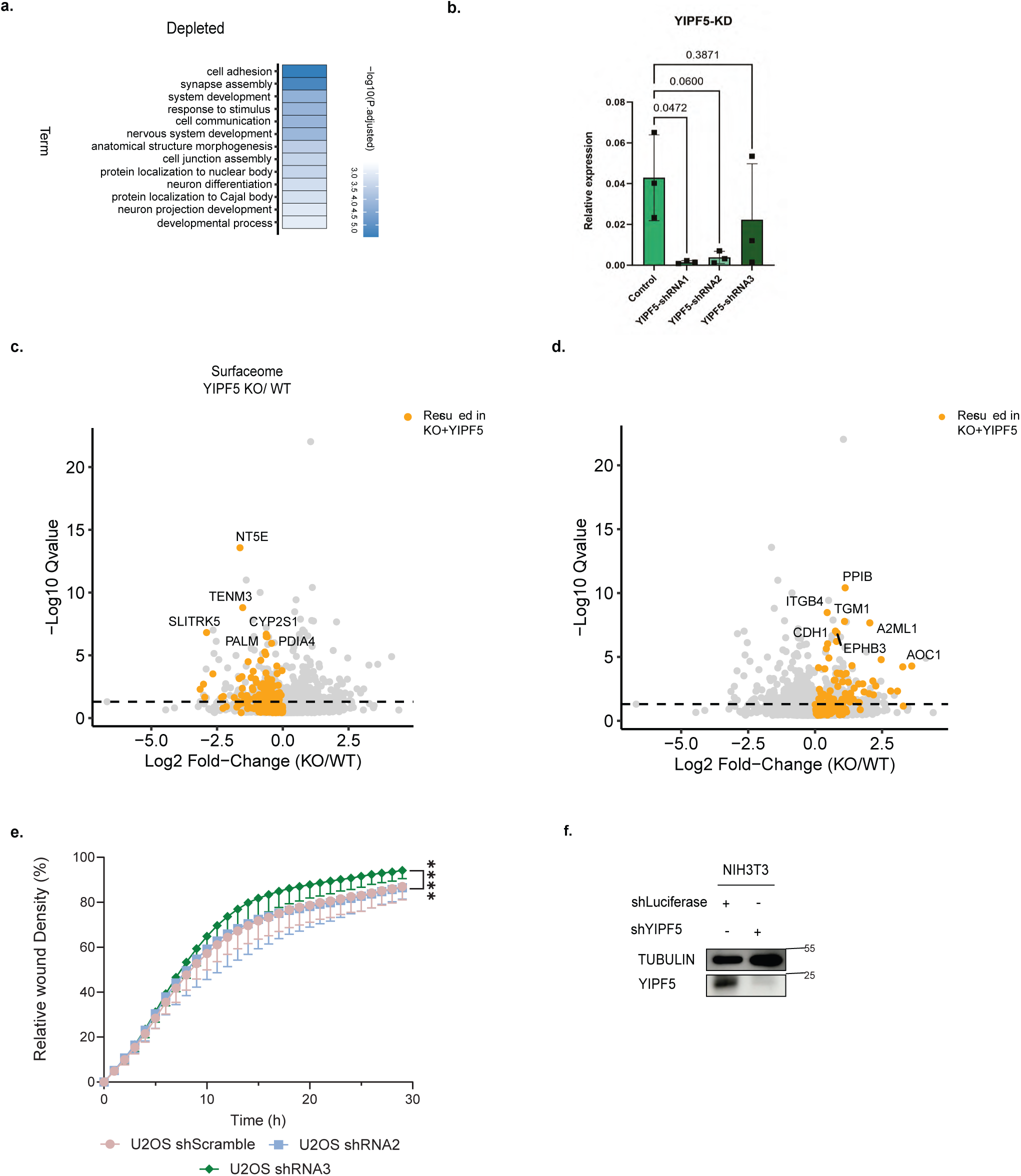
YIPF5 rescues dysregulated protein transport in YIPF5 KO MCF10A cells. Gene Ontology (GO) annotation of proteins with reduced surface expression in YIPF5 KO MCF10A cells compared to WT. Statistically significant categories are represented in shades of blue, with darker blue indicating higher levels of significance. (b) Quantitative PCR analysis of YIPF5 mRNA levels in rat cortical neurons transduced with lentiviruses expressing three distinct shRNAs targeting YIPF5, compared to uninfected controls (n = 3). Statistical significance was determined using a One-way ANOVA with Dunnett’s multiple comparisons test. (c, d) Volcano plots illustrating the rescue of protein transport to the cell surface upon YIPF5 re-expression in YIPF5 KO cells. These plots depict differential surface protein abundance in YIPF5 KO cells re-expressing YIPF5 compared to YIPF5 KO cells. Proteins significantly restored by YIPF5 re-expression are highlighted in yellow, those previously depleted on the surface are shown in (c), those previously accumulated on the surface in (d). The vertical axis represents -log10(q-value), and the horizontal axis represents log2(fold change). (e) U2OS cells stably expressing shScramble, shRNA2 or shRNA3 were plated in Incucyte® Imagelock 96-well plates and a wound healing assay was performed using the Incucyte® Wound Maker 96-Tool on the IncuCyte system with images acquired every hour for 29h. Displayed is the relative wound density at different time points as mean values ± SD from three biological replicates. Statistical significance was evaluated using a two-way ANOVA with Tukey’s multiple comparisons test, with **** denoting p<0.0001. (f) NIH3T3 cells were transfected with plasmids expressing control shLuciferase or shYIPF5 for 72 hours, followed by immunoblotting to detect YIPF5 protein levels, with tubulin as a loading control (n = 3 independent experiments). Source data are available for this figure: SourceData FS4.

**Video 1 YIPF5 depletion induces tubular ERGIC.** Airyscan super-resolution live imaging of MCF10A cells expressing GFP-SURF4 to visualize its localization upon YIPF5 depletion. A WT cell is shown on the left and a YIPF5-KO cell on the right. In YIPF5-KO cells, arrowheads indicate dynamic tubular ERGIC structures. Scale bar = 2 µm. Acquisition rate 517 ms/frame, display frame rate 1,9 frame/second.

**Video 2 Tubular SURF4 structures originate from Sec24C-labeled ERES.** Airyscan super-resolution live imaging of MCF10A YIPF5-KO cells stably expressing GFP-SURF4 and transiently expressing Sec24C-mCherry. The time-stamped video shows the association between SURF4-positive tubules and Sec24C, indicating that tubular ERGIC structures emerge from ERES. Scale bar = 2 µm. Acquisition rate 351 ms/frame, display frame rate 2,8 frame/second.

**Video 3 Tubular SURF4 structures are Rab1b-positive.** Time-stamped Airyscan super-resolution live imaging of MCF10A YIPF5-KO cells stably expressing GFP-SURF4 and transiently expressing Rab1b-mCherry. The video shows colocalization between SURF4-tubules and Rab1b-mCherry, indicating that these dynamic structures are Rab1b-positive. Scale bar = 2 µm. Acquisition rate 1850 ms/frame, display frame rate 0,54 frame/second.

